# Neuronal mimicry generates an ecosystem critical for brain metastatic growth of SCLC

**DOI:** 10.1101/2021.08.10.455426

**Authors:** Fangfei Qu, Siqi Cao, Wojciech Michno, Chioma J. Madubata, Alyssa Puno, Alexandros P. Drainas, Myung Chang Lee, Dian Yang, Angus Toland, Christina Kong, Millie Das, Monte M. Winslow, Anca M. Paşca, Julien Sage

## Abstract

Brain metastasis is a major cause of morbidity and mortality in cancer patients. Here we investigated mechanisms allowing small-cell lung cancer (SCLC) cells to grow in the brain. We show that SCLC cells undergo a cell state transition towards neuronal differentiation during tumor progression and metastasis, and that this neuronal mimicry is critical for SCLC growth in the brain. Mechanistically, SCLC cells re-activate astrocytes, which in turn promote SCLC growth by secreting neuronal pro-survival factors such as SERPINE1. We further identify Reelin, a molecule important in brain development, as a factor secreted by SCLC cells to recruit astrocytes to brain metastases in mice. This recruitment of astrocytes by SCLC was recapitulated in assembloids between SCLC aggregates and human cortical spheroids. Thus, SCLC brain metastases grow by co-opting mechanisms involved in reciprocal neuron-astrocyte interactions during development. Targeting such developmental programs activated in this cancer ecosystem may help treat brain metastases.

## INTRODUCTION

The ability of cancer cells to metastasize to distant sites is a major cause of morbidity and mortality in cancer patients (Chaffer and Weinberg, 2011). In particular, brain metastases are much more frequent than primary brain tumors and occur in 30-40% of all cancer patients, including patients with primary lung tumors, breast tumors, and melanoma (Suh et al., 2020; Valiente et al., 2018). The highly specialized microenvironment in which brain metastases grow puts a unique selective pressure on cancer cells to survive and divide (Boire et al., 2020; Fidler et al., 2010). Clinical advances in treating patients with brain metastases may be aided by a better understanding of the mechanisms that allow cancer cells to grow in this unique microenvironment. Some progress has been made in our understanding of the brain tropism of breast cancer cells (Bos et al., 2009; Ebright et al., 2020; Lorger and Felding-Habermann, 2010; Zeng et al., 2019). Overall, however, the molecular and cellular mechanisms underlying the growth of brain metastases, including from lung cancer, remain a largely unsolved puzzle (Achrol et al., 2019).

Small cell lung cancer (SCLC) is a highly metastatic neuroendocrine carcinoma that accounts for ∼15% of all lung cancers and at least 200,000 deaths worldwide every year (Rudin et al., 2021). Brain metastases are particularly frequent in SCLC patients (Boire et al., 2020). Up to 15% of patients with SCLC have asymptomatic brain metastases at the time of the first diagnosis, and the incidence increases to 40-60% during the course of the disease (Ko et al., 2021; Megyesfalvi et al., 2021; Rusthoven et al., 2020). Brain metastases contribute so much to the morbidity and mortality of SCLC patients that prophylactic cranial irradiation protocols have been implemented, as even slowing the growth of brain metastases might lead to clinical benefit (Neal et al., 2011; Takahashi et al., 2017). Still, the prognosis and overall survival of SCLC patients with brain metastases remain extremely poor (Drapkin and Rudin, 2020; Lukas et al., 2017).

Most efforts to investigate the biology of SCLC brain metastases have focused on the interactions between SCLC cells and blood vessels, as well as how SCLC seed micro-metastases to the brain (Ilhan-Mutlu et al., 2016; Li et al., 2013; Thompson et al., 2013; Xu et al., 2019). However, while the ability of SCLC to expand in the brain is a critical aspect of brain metastases, very little is known about the cellular interactions and molecular programs that drive SCLC growth in the brain microenvironment. SCLC primary tumors have a relative paucity of stromal cells (George et al., 2015), but these tumors can generate their own supportive microenvironment, including by the differentiation of neuroendocrine cancer cells towards less/non-neuroendocrine phenotypes (Lim et al., 2017; Sutherland et al., 2011; Williamson et al., 2016). Whether this or other levels of cellular plasticity impact SCLC growth in the brain microenvironment is unknown, and how SCLC cells interact with resident brain cells has not been investigated (Ko et al., 2021). In large part, this is due to few human samples available for analysis (Lukas et al., 2017; Wang and Komiya, 2019). Furthermore, xenograft models rarely metastasize to the brain (Hodgkinson et al., 2014; Stewart et al., 2020) and the incidence of brain metastasis is extremely rare in genetically engineered mouse models of SCLC (Meuwissen et al., 2003).

Here we sought to investigate the mechanisms that enable the growth of SCLC cells in the brain. Using mouse models and human cortical organoids, we uncovered crosstalk between SCLC cells and astrocytes that is critically for metastatic SCLC growth in the brain. Importantly, these reciprocal interactions are based on neuronal features of SCLC cells, and these neuronal features allow SCLC cells to induce the neuron-protective role of astrocytes. These observations uncover a unique cancer ecosystem that recreates that of the developing brain to enable the growth of brain metastases.

## RESULTS

### Intracranial injection of SCLC cells generates metastases in the brain

To investigate how SCLC cells grow in the brain microenvironment, we generated orthotopic allografts by injecting GFP-expressing mouse SCLC cell lines directly into the brain of recipient mice. We injected SCLC cells into the cerebral cortex because ∼40% of SCLC brain metastases are located in the corresponding frontal and temporal lobe in SCLC patients (Guo et al., 2017). We initially chose to use SCLC cells derived from genetically engineered mouse tumors (Denny et al., 2016; Park et al., 2011) to exclude the possibility of receptor-ligand incompatibility between the murine microenvironment and human cancer cells (Zhang et al., 2005). We used immunodeficient NSG recipient mice to focus on interactions between cancer cells and brain cells and limit interactions with immune cells, including possible activation of T cells by neo-epitopes such as GFP. Mice with SCLC brain allografts survived between 22 and 35 days after injection (**Fig. 1A,B**). At 21 days, transplanted N2N1G (derived from a lymph node metastasis) and 16T (derived from a large primary tumor) cells formed hypercellular tumors with stippled chromatin, nuclear molding, high nuclear-to-cytoplasmic ratios, and frequent mitotic figures and apoptosis. These tumors are morphologically indistinguishable human SCLC brain metastases (**Fig. 1C**). Thus, orthotopic growth of mouse SCLC cells in the mouse brain provide a model to study the mechanisms of metastatic SCLC growth in the brain.

**Figure 1.**
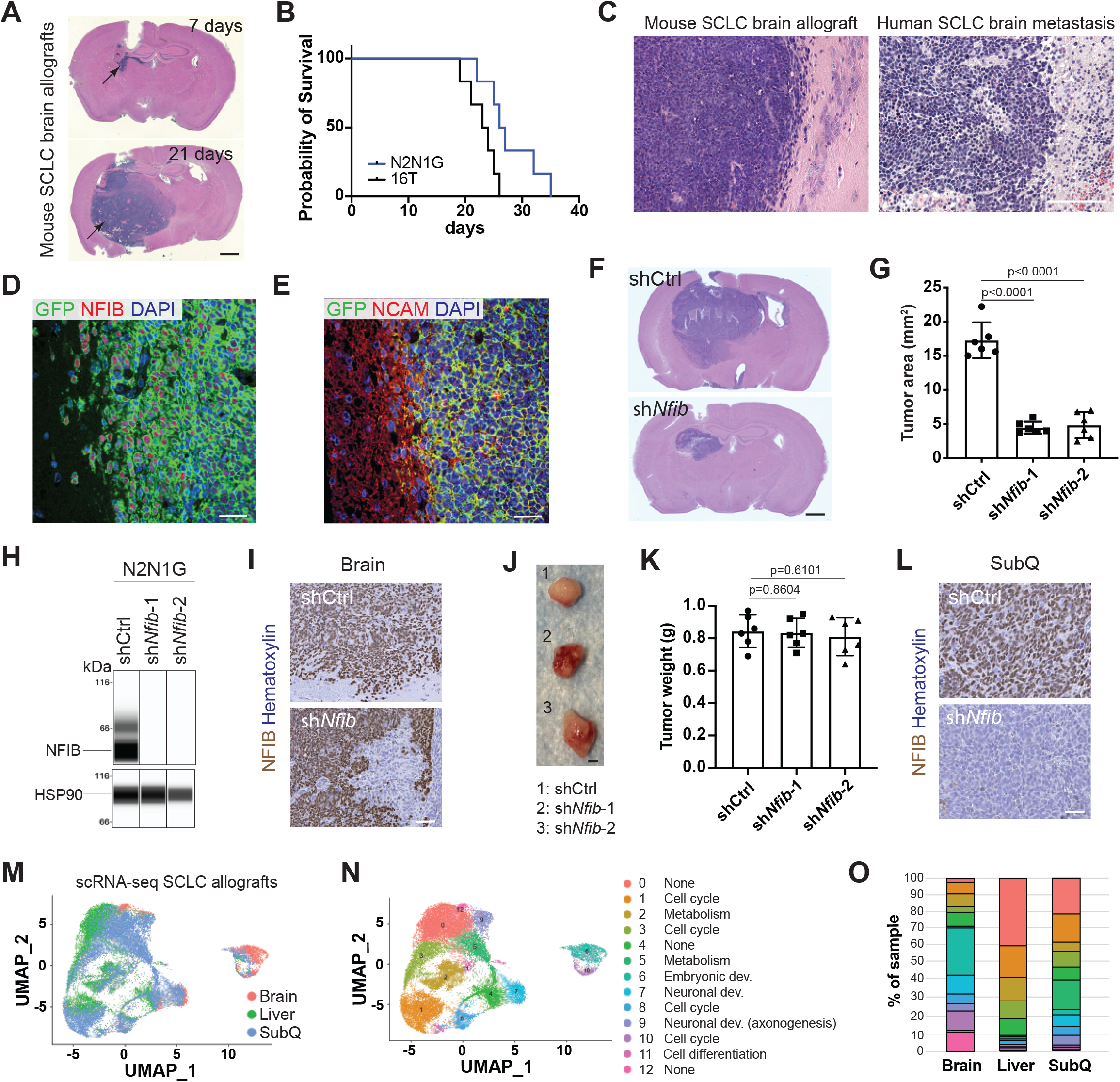
Neuronal features of SCLC cells are associated with their growth in the brain. **A**. Representative images of hematoxylin and eosin (H&E)-stained coronal sections from the brain of mice 7 and 21 days after injection of mouse N2N1G SCLC cells. Arrows show tumors. Scale bar, 1 mm. **B**. Survival plot of mice as in (A) with N2N1G or 16T mouse SCLC cells (n=6 mice each). **C**. Representative comparative H&E-stained sections from brain metastases from mouse N2N1G SCLC cells and from a SCLC patient. Analysis by a certified pathologist (C.K.) confirmed similarities between the two. **D-E**. Representative images of immunofluorescent staining for NFIB (D) and NCAM (E) on sections from SCLC brain allografts (GFP^+^ N2N1G cells). Blue, DAPI to stain DNA. Scale bar, 20 µm. **F-G**. Representative H&E-stained sections and quantification from Control (shCtrl) and sh*Nfib* N2N1G cells 18 days post-injection (n=6 tumors from 2 independent experiments). **H**. Immunoassay (by WES capillary transfer) for NFIB expression in N2N1G cells with control (Ctrl) and *Nfib* shRNAs. HSP90: loading control. **I**. Immunohistochemistry (IHC) for NFIB expression (brown signal) in shCtrl and sh*Nfib* N2N1G brain allograft sections 18 days after transplant. Scale bar, 20 µm. **J-K**. Representative images and quantification of subcutaneous tumors from shCtrl and sh*Nfib* N2N1G cells (n=6 tumors from 2 independent experiments). Scale bar, 5mm. **L**. IHC for NFIB on shCtrl and sh*Nfib* N2N1G subcutaneous tumor sections 18 days after transplant. Scale bar, 20 µm. **M**. UMAP clustering of single-cell gene expression data of N2N1G cells isolated from 2 brain, 3 liver, and 3 subcutaneous tumors. **N**. UMAP clustering plot based on the gene ontology of differentially expressed genes. None indicates no specific gene ontology enrichment for the cluster. **O**. Distribution of cells from brain, liver, and subcutaneous tumors in each cluster (by percentage) (same color legend as in (N)). All p values calculated via two-tailed t-test.

### Neuronal differentiation is associated with the growth of SCLC in the brain

We and others have shown that neuronal programs are upregulated in SCLC cells as tumors progress and become more metastatic (Carney et al., 1982; Denny et al., 2016; Yang et al., 2018, 2019; van Zandwijk et al., 1992). We thus wondered if neuronal differentiation in SCLC contributes to the growth of SCLC in the brain, where neurons functionally interact with non-neuronal cells. As a first test of this idea, we investigated the role of NFIB in SCLC brain tumors. NFIB is a transcription factor frequently up-regulated in late-stage metastatic SCLC (Denny et al., 2016; Dooley et al., 2011; Semenova et al., 2016; Wu et al., 2016). This upregulation drives the expression of neuronal gene programs in SCLC cells and has been implicated in the ability of SCLC cells to migrate using axon-like protrusions, similar to neuroblasts (Denny et al., 2016; Yang et al., 2019). N2N1G and 16T SCLC cells grown in the brain of recipient mice were positive for NFIB and the neuronal marker NCAM (**Fig. 1D,E** and **Fig. S1A**). NFIB knock-down decreased tumor growth in the brain (**Fig. 1F,G** and **Fig. S1B**). Furthermore, brain metastases that formed from NFIB knock-down cells lacked axon-like protrusions (**Fig. S1C,D**). We confirmed initial knock-down of NFIB in N2N1G and 16T cells (**Fig. 1H** and **Fig. S1B**), but the majority of sh*Nfib* N2N1G cells that eventually grew in the brain regained NFIB expression (**Fig. 1I**), suggesting a strong selection pressure to retain NFIB expression in brain metastases. In contrast, NFIB knock-down did not significantly affect tumor growth when cells were injected subcutaneously (**Fig. 1J,K**), as shown previously (Denny et al., 2016), and SCLC cells retained low levels of NFIB (**Fig. 1L**).

In similar orthotopic injection assays, two mouse SCLC cell lines that were derived from primary tumors and express low levels of NFIB failed to consistently grow when injected into the brains of recipient mice (**Fig. S1B,E**) but grew as subcutaneous tumors (**Fig. S1F**). NFIB-high human SCLC cell lines also had a greater ability to form detectable tumors at a frequency higher than 50% in the mouse brain compared to NFIB-low human SCLC cell lines (**Fig. S1E,G**). All these human SCLC cell lines were able to form subcutaneous tumors (**Fig. S1F**).

Collectively, these data indicate that elevated NFIB activity contributes to the ability of SCLC cells to grow in the brain microenvironment. Because NFIB controls neuronal gene programs in SCLC cells (Denny et al., 2016; Yang et al., 2019) and in the developing brain (Fraser et al., 2020; Hickey et al., 2019; Steele-Perkins et al., 2005), these results further suggested that neuronal programs in SCLC cells may contribute to the ability of these cells to grow in the brain microenvironment.

To investigate possible factor in the brain microenvironment influencing SCLC growth in the brain, we performed single-cell RNA sequencing (scRNA-seq) on subcutaneous, liver, and brain allografts. Uniform Manifold Approximation and Projection (UMAP, global and local) analysis uncovered similar cancer cells states across biological replicates; thus, we combined the data from individual tumors to generate organ-specific pooled clusters. (**Fig. S1H-J**). SCLC cells growing in the liver and subcutaneously exhibited similar cancer cell states, suggesting that in these two cases the microenvironment does not have a major influence on transcriptional programs in SCLC cells. In contrast, SCLC cells growing in the brain had distinct gene expression states, suggesting that the brain microenvironment has a unique influence on SCLC cells (**Fig. 1M** and **Fig. S1K,L**). In particular, 42.5% of SCLC cells growing in the brain fell into clusters defined by significantly higher expression (log2FC>1) of genes that have functions in embryonic development and neuronal development comparing to only 3% and 15% of SCLC cells growing in the liver or as subcutaneous tumors, respectively (**Fig. 1N,O**).

Together, these observations suggested that the growth of SCLC cells in the brain may be controlled by both cell-intrinsic and environmental factors. These observations also consolidated our hypothesis that the neuronal features of metastatic SCLC cells may facilitate the growth of brain metastases in part via promoting interactions between SCLC cells and cells in the brain. We thus sought next to identify such functional interactions.

### GFAP-positive reactive astrocytes infiltrate mouse SCLC tumors in the brain

The brain microenvironment constitutes a specialized niche determined in part by brain-specific cellular constituents (Schulz et al., 2019). To determine the brain cell types that may impact SCLC brain metastases, we examined sections from N2N1G and 16T brain allografts for the presence of defined cell types. We found that glial fibrillary acidic protein-positive (GFAP^+^) astrocytes were present within SCLC tumors (**Fig. 2A** and **Fig. S2A**). In contrast, OLIG2^+^ (Oligodendrocyte Transcription Factor 2) oligodendrocytes and MAP2^+^ (Microtubule Associated Protein 2) myelinated neurons were mostly only present at the tumor periphery (**Fig. 2B,C** and **Fig. S2B**). The tumor-associated astrocytes had a hypertrophic morphology with increased cell surface area (**Fig. 2D,E**), similar to reactive astrocytes found around breast cancer brain metastases and in cases of traumatic brain injury (Sirkisoon et al., 2020; Wasilewski et al., 2017).

**Figure 2.**
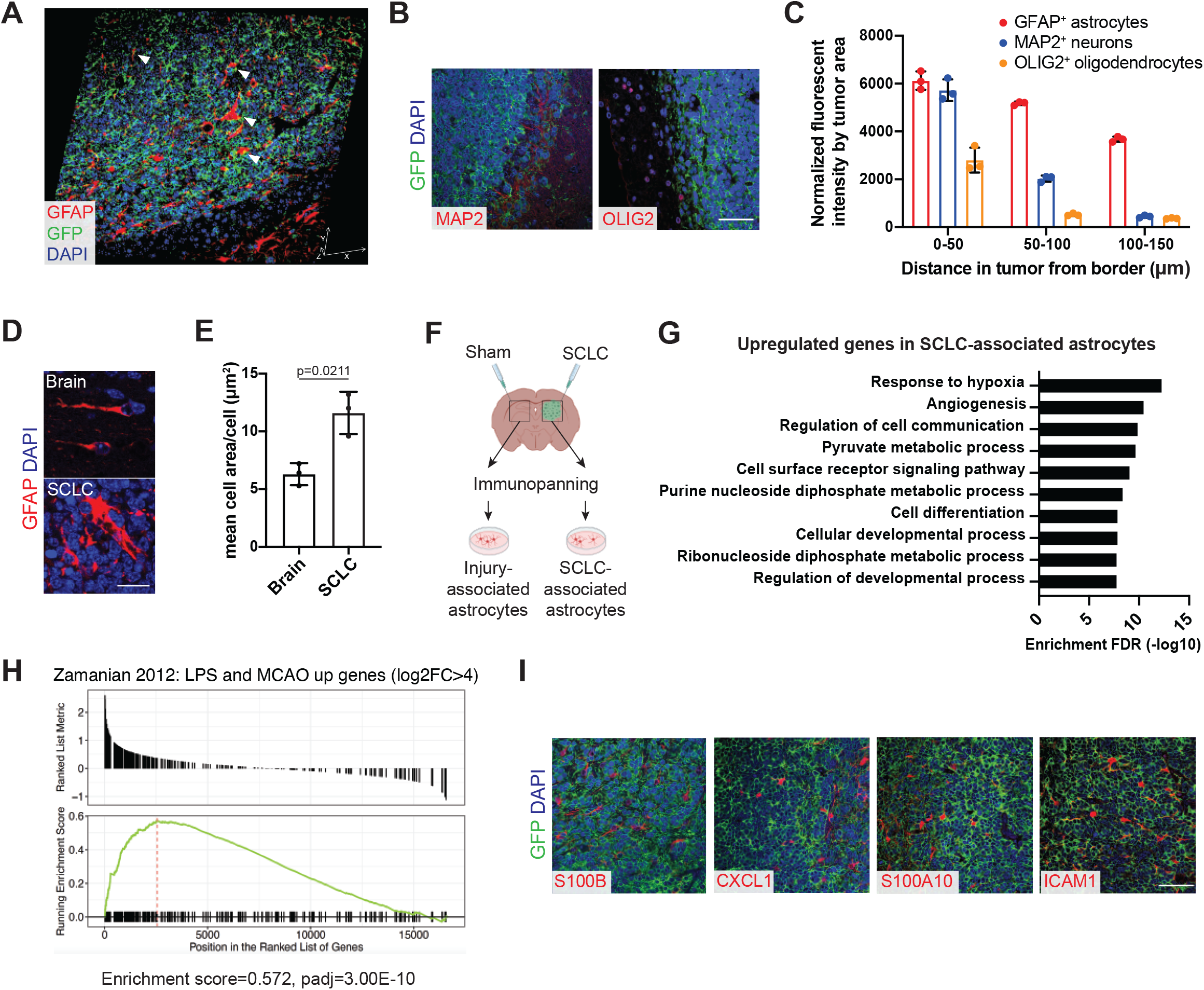
GFAP-positive astrocytes infiltrate mouse SCLC brain metastases. **A**. Representative immunofluorescent staining image of GFAP-positive astrocytes on a section from an N2N1G SCLC brain allograft. Arrowheads show astrocytes. Cancer cells are GFP-positive (green). Blue, DAPI to stain DNA. Scale bars: X, 50 µm, Y, 50 µm, Z, 12 µm. **B**. Representative immunofluorescent staining images of MAP2 (mature neurons) and OLIG2 (oligodendrocytes) on a section from an N2N1G SCLC brain allograft. Cancer cells are GFP-positive (green). Blue, DAPI to stain DNA. Scale bar, 50µm. **C**. Distribution of neurons, astrocytes, and oligodendrocytes as in (A-B) (n=9 tumors from 3 independent experiments, showing means of each experiment). **D**. Representative GFAP-positive (red) astrocyte morphology in the normal adult mouse brain and in an SCLC brain tumor (N2N1G model). Blue, DAPI to stain DNA. Scale bar, 20 µm. **E**. Quantification of cell area (D) (n=3 tumors). P value calculated by two-tailed t-test. **F**. Schematic representation of the immunopanning protocol to isolate SCLC-associated and injury-associated (surgery and sham injection) astrocytes before RNA sequencing and analysis. Astrocytes were isolated from the two sides of the brain of the same mice (N2N1G allograft model). **G**. Gene ontology (GO) enrichment (top 10) for the genes that are upregulated in SCLC-associated astrocytes compared to astrocytes from the sham injection injury site. FDR, false discovery rate. **H**. Gene Set Enrichment Analysis (GSEA) for the genes upregulated in SCLC-associated astrocytes compared to cortical astrocytes from 10-week-old mice for the genes marking reactive astrocytes in endotoxin (LPS) and stroke (MCAO) models. **I**. Representative images of immunofluorescent staining of markers for reactive astrocytes (S100B, CXCL1, S100A10, ICAM1) on sections from N2N1G SCLC brain allografts. Cancer cells are GFP-positive (green). Blue, DAPI to stain DNA. Scale bar, 50µm.

Quiescent astrocytes in the adult brain can be re-activated upon brain injury and regain the ability to proliferate and migrate (Liddelow and Barres, 2017). To better understand the molecular state of GFAP^+^ astrocytes in SCLC brain metastases, we isolated astrocytes from N2N1G brain allografts and contralateral brain regions (following surgery and sham stereotactic injection on this other side) and compared their transcriptome via RNA sequencing (**Fig. 2F**). By principal component analysis (PCA), SCLC-associated astrocytes and sham injection injury-associated astrocytes were more closely related to each other transcriptionally than they were to cortical astrocytes from control young adult mice (no surgery, no injection) (**Fig. S2C**), indicating that the surgery and sham injection were sufficient to influence gene expression programs in astrocytes. However, when we specifically compared SCLC-associated astrocytes and injury-associated astrocytes, we found that SCLC-associated astrocytes upregulate genes involved in cell migration, angiogenesis, hypoxia, and metabolism (**Fig. 2G** and **Fig. S2D**), gene programs previously associated with early brain development and reactive astrogliosis (Cahoy et al., 2008; Henrik Heiland et al., 2019; Zhang et al., 2016; Zou et al., 2019). In addition, SCLC-associated astrocytes showed increased upregulation of genes marking reactive astrocytes from stroke and endotoxin models when compared to wild-type cortical astrocytes (Doron et al., 2019; Liddelow and Barres, 2017; Wang et al., 2020; Zamanian et al., 2012a) (**Fig. 2H, Fig. S2E,F**, and **Table S1**). Immunostaining for various markers of astrocytes reactivation confirmed expression of these factors in SCLC-associated astrocytes (**Fig. 2I**). Thus, astrocytes present in the microenvironment of mouse SCLC brain metastases are reactivated not only by the injury caused by the initial injection of the cancer cells but also by the tumor.

### Human reactive astrocytes infiltrate SCLC tumors

We next addressed whether human astrocytes can also by activated by SCLC tumors. GFAP^+^ astrocytes showed a pattern of tumor infiltration in human SCLC brain metastases that was similar to mouse brain allografts, with deep tumor infiltration (**Fig. 3A**). In contrast, in other cancer types such as breast and lung adenocarcinoma, astrocytes generally surround brain metastasis by forming clear borders without deep infiltration into the tumor lesion ((Priego et al., 2018) and **Fig. 3A**). Indeed, the area occupied by GFAP^+^ astrocytes in tumors was significantly greater in SCLC brain metastases than in the breast cancer, lung adenocarcinoma, and melanoma samples examined (**Fig. 3B**). These observations suggest that astrocytes may interact differently with SCLC tumors than they do with other cancer types.

**Figure 3.**
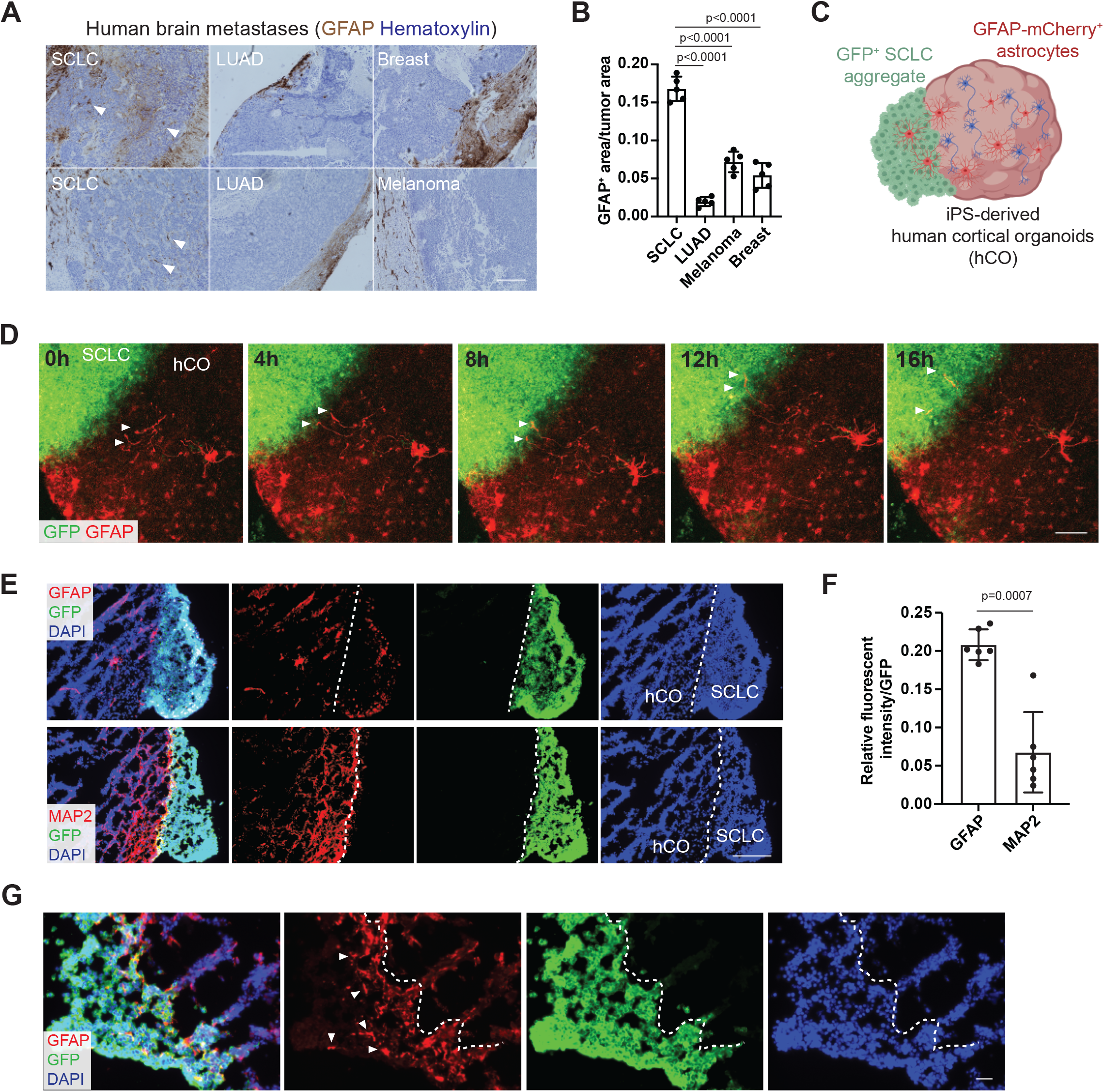
Human GFAP-positive astrocytes infiltrate SCLC. **A**. Representative images showing immunohistochemistry staining for GFAP (brown) on brain metastasis sections counterstained with hematoxylin from SCLC, lung adenocarcinoma (LUAD), breast cancer, and melanoma patients. Arrowheads show astrocytes within SCLC tumors. Scale bar, 500µm. **B**. Quantification of GFAP-positive astrocytes in brain metastases (n=5 tumors from 3 SCLC, 3 LUAD, 3 breast cancer, and 2 melanoma patients). P values calculated via two-tailed t-test. **C**. Schematic description of SCLC-human cortical organoid (hCO) fusion. **D**. Representative images time-series from SCLC-hCO co-cultures showing the astrocyte process movement toward SCLC aggregate at post-fusion day 5-6. Red: GFAP, astrocytes. Green: GFP, SCLC. Scale bar, 100µm. **E**. Representative images showing immunofluorescent staining of GFAP (red), MAP2 (red), and GFP (green for SCLC) on sections from a SCLC-hCO at post-fusion day 10. Blue: DAPI. Scale bar, 200 µm. **F**. Quantification of the relative GFAP or MAP2 fluorescent intensity to GFP in the SCLC aggregates at post-fusion day 10 (n=6 tumors from 2 independent experiments). P value calculated by two-tailed t-test. **G**. Representative image of immunofluorescent staining of GFAP (red) and GFP (green for SCLC) at the SCLC-hCO fusion interface. Blue: DAPI stains for nucleus. Scale bar, 20 µm.

To further determine how human GFAP^+^ astrocytes react to SCLC cells, we generated assembloids by fusion of SCLC aggregates (neuroendocrine SCLC cells normally grow in sphere-like aggregates) with human cortical organoids derived from human induced pluripotent stem (iPS) cells (Paşca et al., 2015; Yoon et al., 2019). After 180-200 days in culture, these organoids contain neurons and glial cells (Paşca et al., 2015; Sloan et al., 2017). To visualize GFAP^+^ astrocytes, we infected day 190 human cortical organoids with an adeno-associated virus (AAV) which express an mCherry fluorescent reporter under the control of the *GFAP* promoter, before co-culturing with SCLC cells at day 200 (**Fig. 3C**). Five days after fusion, we observed active extension of astrocytic processes toward SCLC cells (**Fig. 3D, Fig. S3A**, and **Video S1,2**). Accordingly, at day 10 after fusion, GFAP^+^ astrocytes infiltrated into the SCLC aggregates while MAP2^+^ neurons did not migrate into the SCLC aggregates (**Fig. 3E-G** and **Fig. S3B**). This striking migration of mouse and human GFAP^+^ reactive astrocytes into SCLC tumors identified these cells as potential contributor to a pro-growth SCLC brain ecosystem.

### Paracrine signaling drives functional crosstalk between astrocytes and SCLC cells

To gain a better understanding of the functional interactions between astrocytes and SCLC cells, we developed co-culture assays (**Fig. 4A**). When the two cell types were separated by a membrane, human astrocytes still showed reactive morphology and elevated GFAP expression (**Fig. 4B,C**), consistent with astrocyte activation being mostly driven by a paracrine mechanism. RNA sequencing of SCLC cells co-cultured with human astrocytes showed upregulation of genes that are also found to be upregulated in SCLC cells growing in the brain (**Fig. 1H-J** and **Table S2**), further indicating that this co-culture system recapitulates key aspects of the *in vivo* interactions. Of the genes upregulated in SCLC cells cultured with human astrocytes (log2FC>1, padj<0.05) (**Fig. S4A,B**), more than 20% (18/83) overlap with genes that are specifically upregulated in the SCLC cells from brain metastases (overlap p=9.35e-05) (**Fig. S4C**). Among these conserved genes, nine are intimately related to nervous system development (**Fig. S4C**), in part recapitulating our observations of an induction of neuronal programs in SCLC cells growing in the brain (**Fig. 1H-J**).

**Figure 4.**
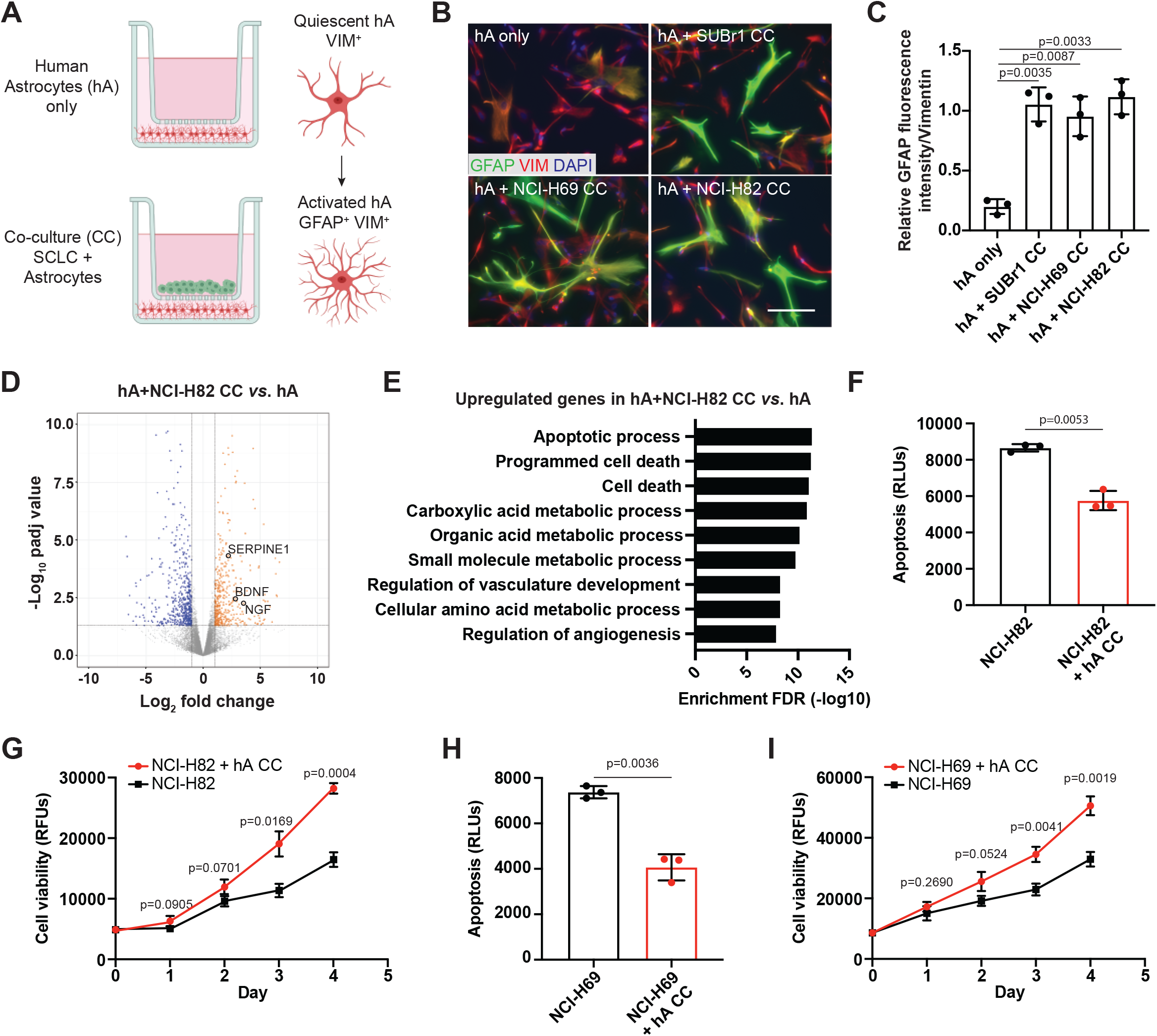
Astrocytes activated by SCLC cells secrete factors promoting the survival of SCLC cells. **A**. Schematic representation of co-culture (CC) assays with human astrocytes (hA) and SCLC cells. **B**. Representative images showing immunofluorescent staining of human astrocytes (hA) cultured alone or co-cultured with human SCLC cell lines (SUBr1, NCI-H69, and NCI-H82). GFAP (green), Vimentin (VIM, red). Blue, DAPI to stain DNA. Scale bar, 20µm. **C**. Quantification of GFAP fluorescent intensity relative to Vimentin (n=6 from 3 independent experiments). **D**. Volcano plot showing differentially expressed genes in hA co-cultured with NCI-H82 cells compared to hA cultured alone. Selected genes are indicated. **E**. Gene ontology (GO) enrichment (top 10) for the genes that are upregulated in hA co-cultured with NCI-H82 cells. **F**. Apoptosis (caspase3/7 activity) measured in NCI-H82 cells cultured with (red) or without (black) hA (n=3 independent experiments). **G**. Cell viability (AlamarBlue assay) measured in NCI-H82 cells cultured with (red) or without (black) hA (n=3 independent experiments). **H-I**. Apoptosis (H) and cell viability (I) in NCI-H69 cells as in (F-G). All p values calculated by two-tailed t-test.

To determine the extent and nature of the changes induced in astrocytes by SCLC cells, we performed RNA sequencing of human astrocytes co-cultured for 5 days with two human SCLC cells (NCI-H82 and NCI-H69) (**Fig. 4D, Fig. S4D**, and **Table S3**). Several genes upregulated in SCLC brain tumor-associated astrocytes, including *ASNS, GDF15, ICAM1*, and *CXCL1*, were also significantly upregulated in human astrocytes co-cultured with NCI-H82 and NCI-H69. Unsupervised clustering analysis showed that human astrocytes co-cultured with human SCLC cells grouped together (**Fig. S4E,F**). Gene ontology (GO) analysis of the upregulated genes in astrocytes co-cultured with SCLC cells highlighted pathways that were also enriched in SCLC-associated astrocytes (e.g., hypoxia, metabolism; see **Fig. 2G** and **Fig. S2D**), along with enrichment in programs involved in cell death (**Fig. 4E, Fig. S4G**, and **Table S4**). In particular, we noticed an upregulation of genes coding for neuroprotective factors that are normally secreted by astrocytes to support the survival of neurons during early brain development or following acute injuries, such as NGF (Nerve Growth factor), BDNF (Brain Derived Neurotrophic Factor), TGF-α (Transforming growth factor alpha), and SERPINE1 (Serpin Family E Member 1) (Almeida et al., 2005; Fang et al., 2012; Nguyen et al., 2009).

These data suggested that astrocytes may support the survival of SCLC cells *via* similar mechanisms to those protecting neurons from apoptosis. Indeed, SCLC cells grew better and showed decreased apoptotic cell death when co-cultured with astrocytes (**Fig. 4F-I** and **Fig. S4H-K**). Notably, conditioned medium from astrocytes that were never in contact with SCLC cells or SCLC-conditioned medium did not have pro-survival effects on SCLC cells (**Fig. S4L**). Thus, crosstalk between SCLC cells and astrocytes is necessary for the activation of astrocytes to provide these pro-survival effects to SCLC cells, further highlighting the extensive functional interactions between SCLC cells and astrocytes.

### Secretion of the neuronal survival factor SERPINE1 by astrocytes supports the growth of SCLC cells

To identify specific pro-survival secretory molecules that are produced by SCLC-activated astrocytes, we compared upregulated genes in human astrocytes cultured with human SCLC cells (**Fig. 3**) and mouse astrocytes associated with SCLC brain allografts (**Fig. 2**). This analysis identified 11 genes whose expression was found to be consistently upregulated (**Fig. 5A**). While it is likely that several of these factors contribute to the pro-survival effects of astrocytes on SCLC cells, we chose to focus on SERPINE1 (also known as Plasminogen Activator Inhibitor 1 or PAI-1), a secretory molecule that inhibits apoptosis in several contexts (Che et al., 2018; Pavón et al., 2016; Schneider et al., 2008). We confirmed that SERPINE1 protein expression is elevated in astrocytes upon co-culture with human SCLC cells (**Fig. 5B,C**). We also noted that SERPINE1 is expressed in GFAP-positive astrocytes associated with mouse SCLC brain tumors (**Fig. 5D,E**) and human SCLC brain metastases (**Fig. 5F**) as well as in the SCLC/human cortical organoids assembloids (**Fig. S5A**). Addition of recombinant SERPINE1 to the culture medium was sufficient to increase the expansion of SCLC cells (**Fig. 5G**). SERPINE1 inhibition by the potent and selective inhibitor Tiplaxtinin (Tip) (Che et al., 2018; Elokdah et al., 2004; Pavón et al., 2016) reduced the expansion of SCLC cells, and this was partially rescued by adding recombinant SERPINE1 (**Fig. 5G,H** and **Fig. S5B**). Moreover, Tip reduced the pro-growth and anti-apoptotic effects of SCLC-activated astrocytes in both human (**Fig. 5H,I** and **Fig. S5C-H**) and mouse cell lines (**Fig. 5J-L**). Treatment of N2N1G SCLC cells with Tip once at the time of intra-cranial injection led to significant tumor inhibition 2 weeks later (**Fig. 5M** and **Fig. S5I**), suggesting that inhibition of SERPINE1 at the time of seeding may have an effect on the long-term growth of brain metastases. Thus, astrocytes activated by SCLC cells promote the expansion of SCLC cells at least in part by promoting cell survival, including through the secretion of SERPINE1.

**Figure 5.**
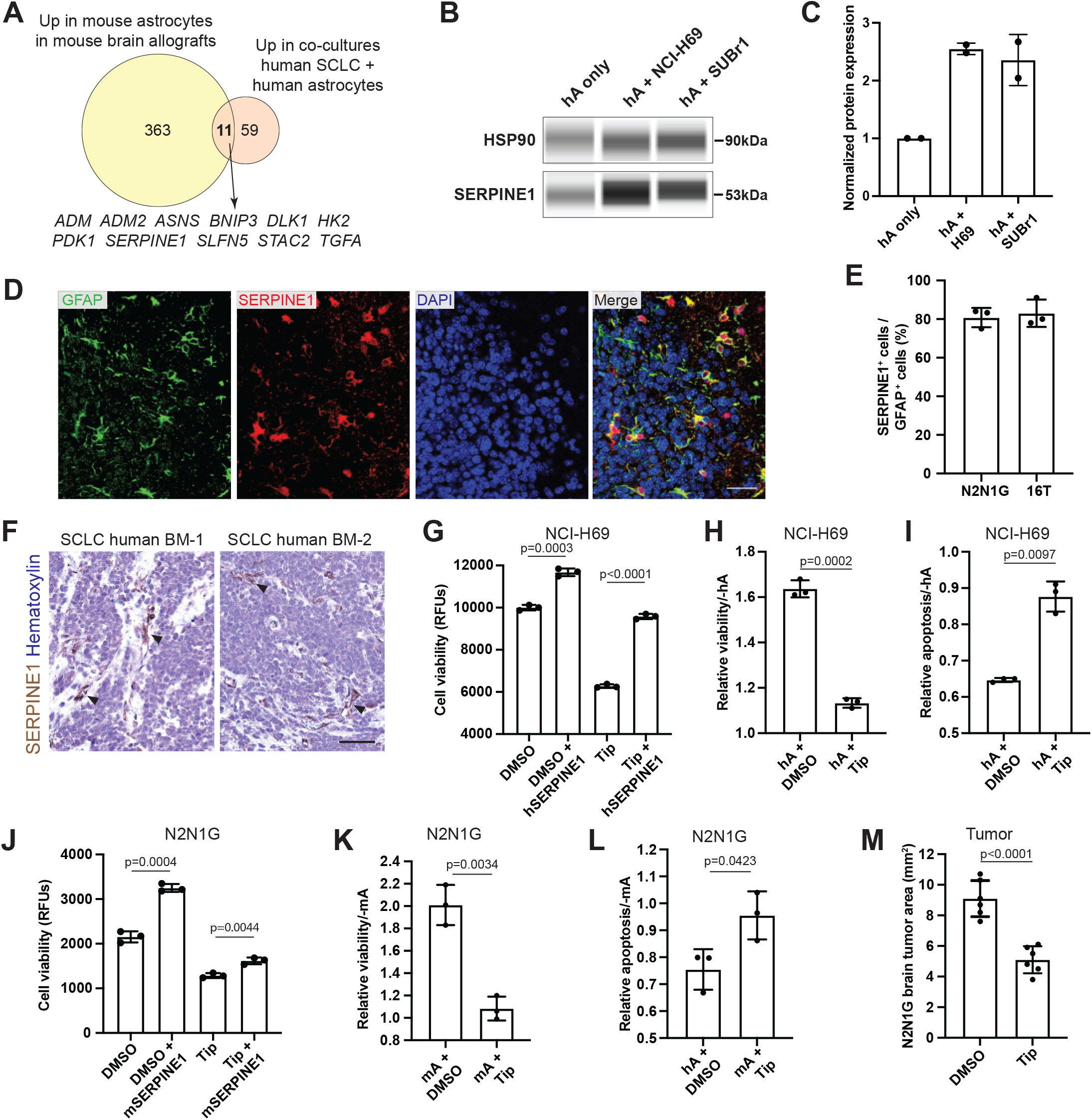
SERPINE1 is upregulated in reactive astrocytes and is required for optimal SCLC growth. **A**. Overlap between upregulated genes in tumor-associated mouse astrocytes (mA) and human astrocytes (hA) co-cultured with SCLC cells. The 11 overlapping genes are indicated. **B**. Immunoassay (by WES capillary transfer) for SERPINE1 expression in naive hA and hA co-cultured with the NCI-H69 and SUBr1 human SCLC cell lines. HSP90 serves as a loading control. **C**. Quantification of (B) (n=2). **D**. Representative immunofluorescent staining images of GFAP (green) and SERPINE1 (red) in a brain tumor from N2N1G mouse SCLC cells. Blue, DAPI to stain DNA. Scale bar, 50 µm. **E**. Quantification of the overlap between SERPINE1 and GFAP from images as in (D) in two SCLC allograft models (N2N1G and 16TG, n=3 tumors each). **F**. Representative images showing immunohistochemistry for SERPINE1 (brown, arrowhead) on SCLC brain metastasis sections. Scale bar, 50 µm. **G**. Cell viability (AlamarBlue assay) measured in NCI-H69 cells treated with recombinant hSERPINE1 and the SERPINE1 inhibitor Tiplaxtinin (Tip) (n=3 independent experiments). **H-I**. Cell viability (AlamarBlue assay) and apoptosis (caspase3/7 activity) measured in NCI-H69 cells cultured with hA compared to without hA, with or without Tip treatment (n=3 independent experiments). **J**. Cell viability (AlamarBlue assay) measured in N2N1G cells treated with recombinant mSERPINE1 and Tip (n=3 independent experiments). **K-L**. Cell viability (AlamarBlue assay) and apoptosis (caspase3/7 activity) measured in N2N1G cells cultured with mA compared to without mA, with or without Tip treatment (n=3 independent experiments). **M**. Quantification of tumor area after intra-cranial injection of N2N1G mouse cells with DMSO control or Tip (n=6 tumors from 2 independent experiments). All p values calculated by two-tailed t-test.

### Reelin-VLDLR signaling is required for astrocyte chemotaxis induced by SCLC cells

To identify signals from SCLC cells that induce migration and/or activation of astrocytes, we next examined receptor-ligand pairs for genes that are upregulated in tumor-associated astrocytes (**Fig. 2**) and neuronal genes that are upregulated (log2FC>1, padj<0.05) in metastatic SCLC cells (from (Yang et al., 2019), see also **Table S5**). Using this approach, we identified the Reelin-VLDLR pair as a candidate mediator of functional interactions between astrocytes and SCLC cells in the brain, with upregulation of Reelin in metastatic SCLC and upregulation of its receptor VLDLR in tumor-associated astrocytes (**Fig. 6A**). We confirmed that Reelin was expressed in N2N1G SCLC cells growing in the brain (**Fig. 6B,C** and **Fig. S6A**) and human SCLC brain metastases (**Fig. 6D**). Also, VLDLR was expressed in brain tumor-associated astrocytes (**Fig. 6B,C** and **Fig. S6B**). When we co-cultured human astrocytes and shControl or sh*Reln* NCI-H69 cells and assessed levels of GFAP (marking re-activated astrocytes) and Vimentin, we found that sh*Reln* NCI-H69 cells could still induce GFAP expression in human astrocytes comparable to control cells (**Fig. 6E-G**). Astrocytes could also still support SCLC growth when Reelin was knocked-down in SCLC cells (**Fig. S6C**). Moreover, recombinant human Reelin did not elevate GFAP expression in human astrocytes (**Fig. S6D**). Thus, Reelin is dispensable for SCLC-mediated astrocyte activation.

**Figure 6.**
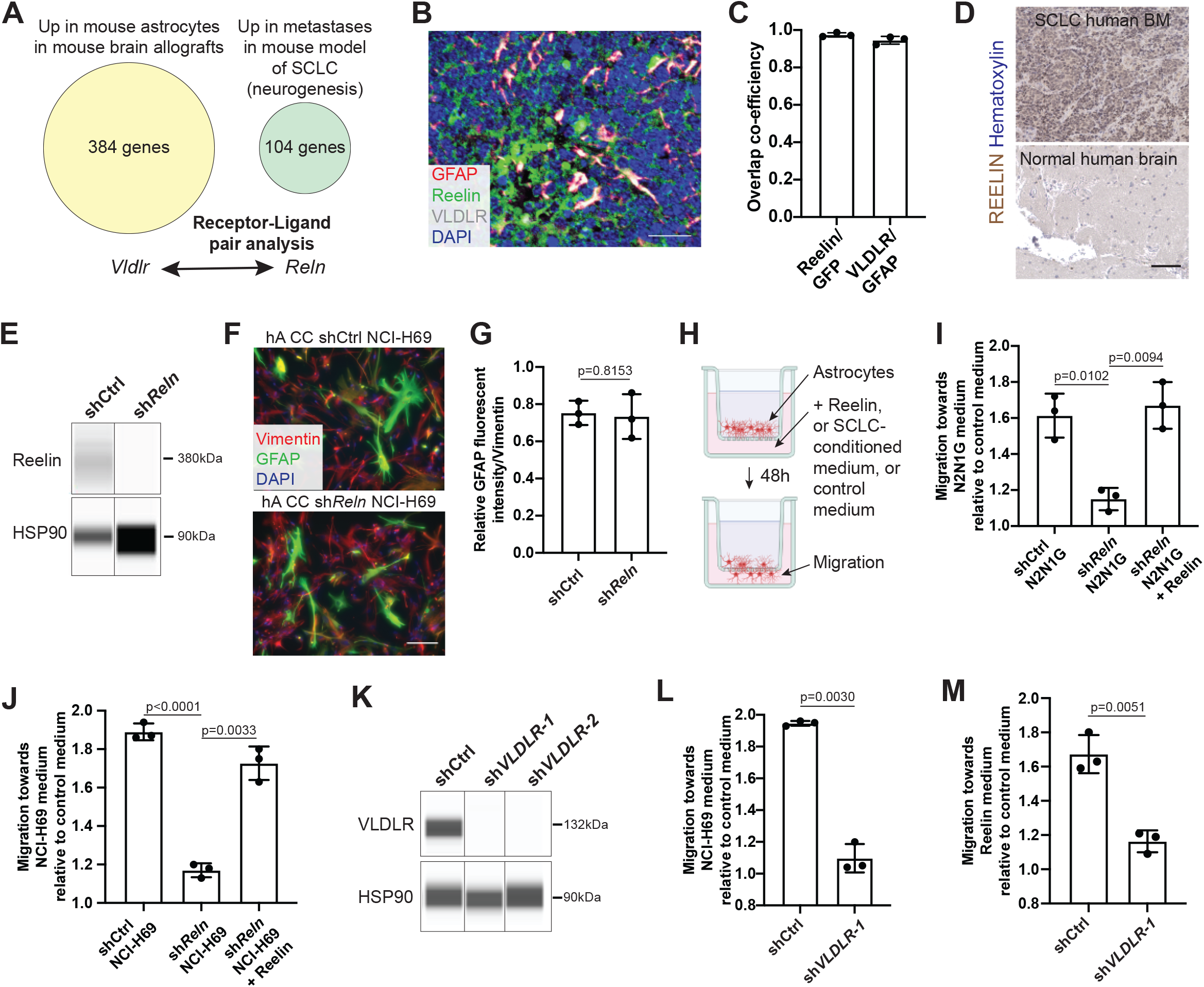
Reelin-VLDLR signaling regulates SCLC-induced astrocyte migration. **A**. Receptor-ligand pair analysis on upregulated neuronal genes in SCLC metastases and upregulated genes in the SCLC-associated astrocytes. **B**. Representative image of immunofluorescence staining of Reelin (green), VLDLR (gray) and GFAP (red) on an N2N1G brain allograft section. Blue: DAPI to stain DNA. Scale bar, 20 µm. **C**. Quantification of co-localization of Reelin with GFP (which labels SCLC cells) and VLDLR with GFAP (which labels astrocytes) in N2N1G brain allografts (n=3). **D**. Representative images showing immunohistochemistry for Reelin (brown) on sections from human SCLC brain metastases and normal brain regions. Scale bar, 100 µm. **E**. Immunoassay (by WES capillary transfer) for Reelin expression in shControl and sh*Reln* N2N1G cells. HSP90: loading control. **F**. Representative images showing immunofluorescent staining of GFAP (green) and Vimentin (red) in human astrocytes (hA) co-cultured (CC) with shControl or sh*Reln* NCI-H69 SCLC cells. Blue: DAPI to stain DNA. Scale bar, 20 µm. **G**. Quantification of fluorescent intensity GFAP normalized to Vimentin (n=3). **H**. Astrocyte chemotaxis assay in the presence or absence of SCLC-conditioned culture medium. Migration is measured by fluorescent intensity (f.i.) of live cells in the bottom well. **I-J**. Chemotaxis of astrocytes in shControl and sh*Reln* N2N1G (H, mA) and NCI-H69 (I, hA)-conditioned culture medium compared to control culture medium as in (H), with or without addition of recombinant Reelin (n=3 independent experiments). **K**. Immunoassay (by WES capillary transfer) for VLDLR expression in shControl and sh*VLDLR* hA. **L-M**. Chemotaxis of shControl and sh*VLDLR*-1 hA in NCI-H69-conditioned culture medium (L) and culture medium containing recombinant human Reelin protein (M) compared to control culture medium. All p values are calculated via two-tailed t-test.

Reelin-VLDLR signaling has a critical function in regulating neuronal and glial progenitor migration during early development (Brunkhorst et al., 2015; Courtès et al., 2011; Zhan et al., 2017). Therefore, we hypothesized that elevated Reelin expression by metastatic SCLC could cells promote the migration/chemotaxis of reactivated astrocytes with higher VLDLR level. To test this idea, we established a Transwell-based chemotaxis assays where astrocytes were allowed to migrate toward medium containing recombinant Reelin, SCLC-conditioned medium, or control medium (**Fig. 6H**). In this assay, recombinant mouse Reelin was sufficient to promote the migration of mouse astrocytes, and migration towards SCLC-conditioned medium was reduced by a Reelin-blocking antibody (**Fig. S6E,F**). Conditioned medium from mouse N2N1G SCLC cells also increased the migration of astrocytes in culture which could be prevented by Reelin knock-down in the cancer cells (**Fig. 6I** and **Fig. S6G**). Similar observations were made with human SCLC cells and human astrocytes. Human SCLC-conditioned medium and human recombinant Reelin protein increased migration of human astrocytes (**Fig. S6H,I**) and such pro-migration effect is lost in medium conditioned with sh*Reln* human SCLC cells (NCI-H69) (**Fig. 6J** and **Fig. S6J**). Knock-down of the Reelin receptor VLDLR in primary human astrocytes significantly inhibited human astrocyte migration (**Fig. S6K**) as well as chemotaxis toward SCLC-conditioned medium or medium with recombinant Reelin (**Fig. 6L,M** and **Fig. S6L-N**). Collectively, these results indicate that the migration of astrocytes towards SCLC cells is controlled at least in part by Reelin produced by SCLC cells and its receptor VLDLR expressed by astrocytes.

### Reelin is specifically required for SCLC growth in the brain

Knockdown of Reelin did not affect the expansion of N2N1G and 16TG cells *in vitro* and did not change levels of apoptosis under normal culture conditions (**Fig. S7A,B**), indicating that Reelin is not intrinsically required for the growth of SCLC in this setting. To determine if Reelin expression in SCLC cells contributes to tumor growth *in vivo*, we transplanted sh*Reln* and control N2N1G and 16T cells into recipient mice. We found that Reelin knock down in N2N1G and 16T cells had no effect on subcutaneous tumor growth (**Fig. 7A,B** and **Fig. S7C**) or on the growth of liver metastases after intravenous injection (**Fig. 7C-E** and **Fig. S7D**). In contrast, N2N1G and 16T cells with Reelin knockdown generated smaller tumors in the brain compared to controls (**Fig. 7F,G** and **Fig. S7E**). Reelin knockdown N2N1G cells growing in these three sites retained low expression of Reelin (**Fig. S7F**). Astrocyte infiltration was significantly reduced in the mouse brain metastases generated from sh*Reln* N2N1G cells compared to controls (**Fig. 7H,I**). The migration of sh*Reln* N2N1G cells themselves was not affected in a 3D Matrigel model (**Fig. S7G,H**). In the brain, tumors generated by sh*Reln* N2N1G and 16TG cells showed significantly increased apoptosis compared to control cells (**Fig. 7J,K** and **Fig. S7I**). Together, these experiments indicate Reelin produced by SCLC cells is required for the growth of SCLC tumors specifically in the brain microenvironment.

**Figure 7.**
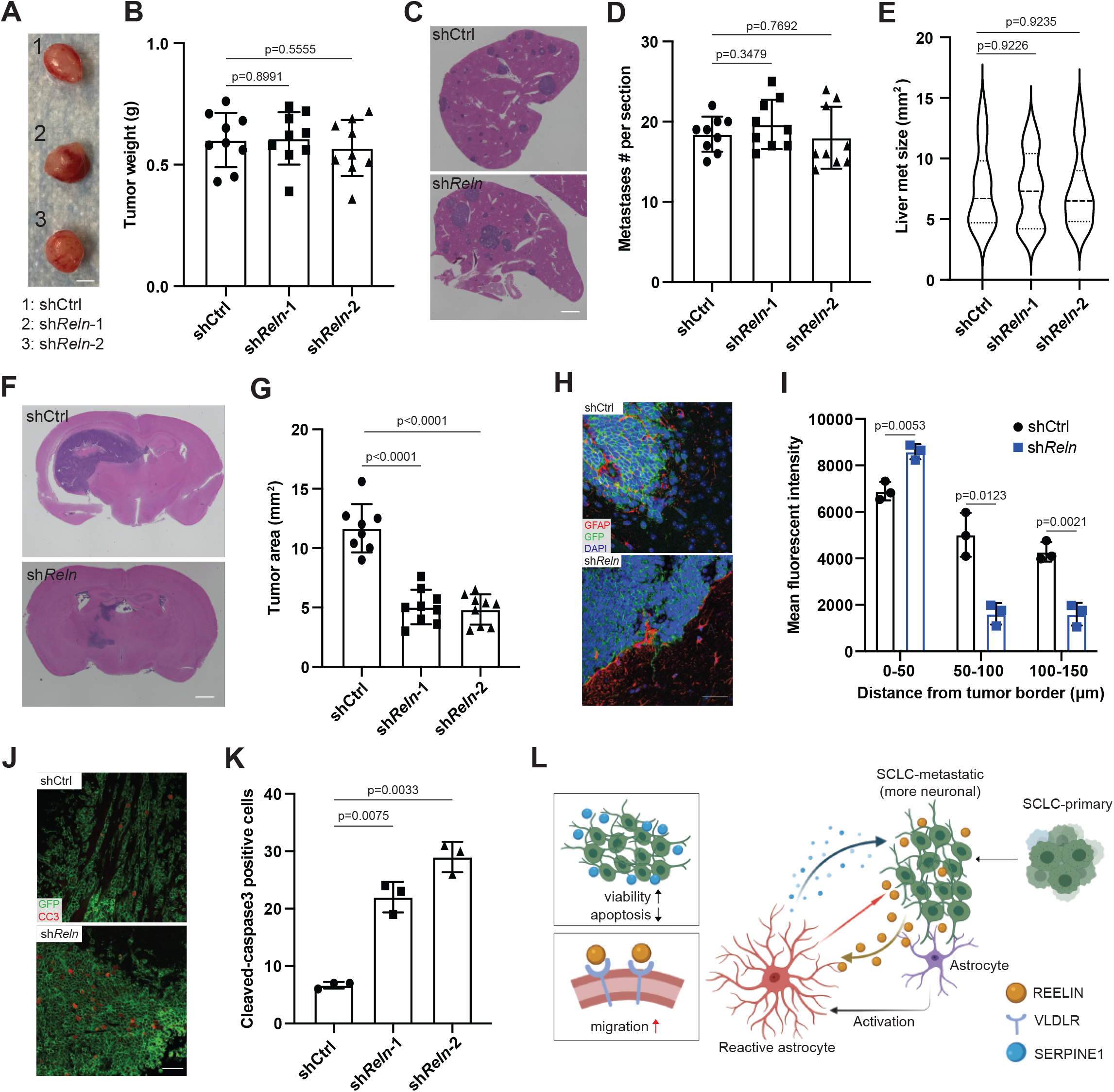
Reelin expression is critical for the growth of SCLC brain tumor *in vivo*. **A**. Representative images of subcutaneous tumors generated by shControl N2N1G cells and sh*Reln* N2N1G cells. **B**. Quantification of the size of subcutaneous tumors as in (A). N=9 from 2 independent experiments. Scale bar, 5mm. **C**. Representative images of metastases on liver sections (H&E stained) generated from i.v. injection of shControl and sh*Reln* N2N1G cells. **D**. Number of liver metastases generated as in (C). N=9 from 2 independent experiments. Scale bar, 2mm. **E**. Tumor size of liver metastases of (C). N=15 tumors analyzed from each group. **F**. Representative images of shControl and sh*Reln* N2N1G brain allografts 21 days post-transplant. Purple: tumor. Scale bar, 2mm. **G**. Tumor size of brain allografts as in (E). N=8, 9 mice from 3 independent experiments. **H**. Representative images of immunofluorescent staining of GFAP (red) on SCLC brain tumors generated by shControl and sh*Reln* N2N1G cells (green). Scale bar, 20µm. **I**. Astrocyte infiltration in brain allografts quantified by the mean fluorescent intensity divided by the distance from tumor border. Black dot: shControl N2N1G. Blue square: sh*Reln* N2N1G. **J**. Representative images of immunofluorescent staining of cleaved caspase3 (CC3, red) on the shControl and sh*Reln* N2N1G cells (green) growing in the brain of mice. Scale bar, 20µm. **K**. Quantification of the number of CC3-positive N2N1G cells as in (I). N=8 mice from 3 independent experiments. All p values are calculated via two-tailed t-test. **L**. Working model: SCLC brain metastases show upregulation of neuronal features compared to primary tumors and these neuronal features play a critical role in the ecosystem between SCLC cells and astrocytes. Secretion of Reelin by SCLC cells recruit re-activated astrocytes to the tumor site. Secretion of SERPINE1 by astrocytes in turn promote the survival of SCLC cells.

## DISCUSSION

While brain metastases are very common in SCLC patients and cause high morbidity and mortality, little is known about how SCLC cells survive and grow into tissue-destructive metastases in the brain (Drapkin and Rudin, 2020; Lukas et al., 2017; Wang and Komiya, 2019). Here we investigated the crosstalk between SCLC cells and cells in the brain microenvironment, focusing on how neuronal programs in SCLC cells contribute to their growth in the brain. We uncovered reciprocal interactions between SCLC cells and reactive astrocytes (**Fig. 7L**). Strikingly, the mechanisms underlying these interactions are similar to those that occur between neurons and astrocyte progenitors during brain development. This co-optation of brain development mechanisms by SCLC cells could open new therapeutic avenues to prevent or treat brain metastases.

The brain constitutes a unique microenvironment that cancer cells must adapt to as they growth into metastases. Brain cells could serve as a supportive stroma for cancer growth by producing generic pro-cancer signals (e.g. pro-survival, or pro-proliferation), but emerging evidence suggests that cancer cells across several cancer types can acquire neuronal features during tumor progression, which may endow them with the capacity to interact more specifically with cells in the brain (Caino et al., 2016; Denny et al., 2016; Neman et al., 2014; Tan et al., 2020; Wingrove et al., 2019). There is also evidence of transcriptomic changes within brain cells when tumors are present (Quail and Joyce, 2013; Sevenich et al., 2014; Wingrove et al., 2019). However, our understanding of how specific neuronal genes/programs facilitate the crosstalk between cancer cells and the brain microenvironment to benefit tumor growth remains limited. Our work shows that astrocytes can be re-activated by SCLC cells *via* paracrine mechanisms, although the identity of the specific factors secreted by SCLC cells remains unknown. We further identify a number of molecules produced by both these reactivated astrocytes and SCLC cells as the two cell types interact. In particular, we uncovered a role for Reelin produced by SCLC cells and its receptor VLDLR on the surface of astrocytes in SCLC growth in the brain. We note that VLDLR is also expressed by SCLC and our previous work showed that VLDLR knock-down can lead to a slight decrease in the ability of SCLC cells themselves to migrate in culture (Yang et al., 2019). It is likely that other functional interactions remain to be discovered but our data provide a clear indication of autocrine and paracrine mechanisms between cancer cells and cells in the brain during the metastatic process. In this context, cortical spheroids derived from iPS cells are a powerful tool to study cell-cell interactions in the brain (Choe et al., 2020; Paşca et al., 2015) and may provide a relevant platform to investigate how neuronal programs in SCLC cells enhance the ability of these cells to grow in the brain.

SCLC tumors often have few non-cancer cells (∼85% cancer cells in primary tumors) (George et al., 2015). We and others have previously shown that SCLC cells that have differentiated into less/non-neuroendocrine states can serve as a supportive stromal-like population (Calbo et al., 2011; Hodgkinson et al., 2014; Lim et al., 2017; Pan et al., 2021; Williamson et al., 2016). Notably, non-neuroendocrine cells generated by activation of Notch within tumors seem to be less frequent or even absent in metastases (Lim et al., 2017), possibly because of a lack of signals to activate Notch signaling, which raises the question whether other cell types in the metastatic ecosystem may functionally replace these non-neuroendocrine cancer cells. Intriguingly, genes that are upregulated in these supportive Notch-positive non-neuroendocrine SCLC cells showed enrichment for an astroglial signature (Lim et al., 2017). Midkine, a protein normally produced by astrocytes to promote brain development, is expressed Notch-positive non-neuroendocrine SCLC cells and can promote the survival of neuroendocrine SCLC cells (Lim et al., 2017). SERPINE1 is also upregulated in these non-neuroendocrine SCLC cells (Lim et al., 2017). Therefore, it is possible that neuroendocrine SCLC cells are intrinsically poised to benefit from factors secreted by astrocytes, which could in part explain the frequent growth of SCLC brain metastases in patients.

Even though few GFAP^+^ astrocytes penetrate inside breast cancer, lung adenocarcinoma, and melanoma brain metastases, these GFAP^+^ astrocytes have been shown to play pro-tumorigenic roles in these tumors, including inducing angiogenesis and forming gap-junctions for nutrient support for cancer cells at the tumor border (Chen et al., 2016; Kim et al., 2011; Priego et al., 2018; Wasilewski et al., 2017). SCLC cells may also share some of these abilities with other cancer types to benefit from the presence of reactive astrocytes. In our *in vitro* assays, we used a Transwell system for non-contact co-culture but it is likely that the physical contact between SCLC cells and astrocytes also play important roles in their crosstalk. In addition, the activity of tumor-associated astrocytes may affect the immune response in the brain microenvironment, including microglia and macrophages (Berghoff et al., 2016; Henrik Heiland et al., 2019; Hohensee et al., 2017; Sirkisoon et al., 2020). Thus, it will be important for future studies to determine the systematic changes in various immune components within the SCLC brain metastases using immunocompetent models.

Therapeutically, targeting the pathways involved in the functional interactions between SCLC cells and astrocytes may benefit SCLC patients, either patients with brain metastases or as a prophylactic measure similar to cranial irradiation. For instance, SERPINE1 inhibitors have been investigated in several contexts already, including cancer (Pavón et al., 2016; Placencio and DeClerck, 2015), fibrosis (Brown, 2010; Lee et al., 2012), and Alzheimer’s disease (Jacobsen et al., 2008; Kutz et al., 2012). There are no specific inhibitors for Reelin and VLDLR, but several studies have indicated the therapeutic potential of targeting Reelin and Reelin signaling pathways in different cancer types (Jandial et al., 2017; Lin et al., 2016; Qin et al., 2017). In the future, treating brain metastases by targeting pathways specifically implicated in brain development may minimize side effects in the adult brain.

## ACKNOWLEDGEMENTS

We thank all the members of the Sage lab for their help and support throughout this study, including Andy He for his help with the isolation of SUBr1 cells, and Pauline Chu for her help with tissue sections. Research reported in this publication was supported by the NIH (J.S. CA231997 and CA217450; C.K. CA231997), a Damon Runyon Cancer Research Foundation fellowship (F.Q.), Stanford Graduate Fellowships (D.Y., M.C.L.), and a TRDRP dissertation award (D.Y.). J.S. is the Harriet and Mary Zelencik Scientist in Children’s Cancer and Blood Diseases and the Elaine and John Chambers Professor in Pediatric Cancer.

## AUTHOR CONTRIBUTIONS

Conceptualization: F.Q., M.M.W., A.M.P., J.S.

Methodology: F.Q., D.Y.

Formal Analysis: F.Q., S.C., W.M., C.J.M., A.P.D.

Investigation: F.Q., S.Q., W.M., A.P., M.C.L., A.T., C.K., M.D.

Writing – Original Draft: F.Q., J.S.

Writing – Reviewing and Editing: all authors

Visualization: F.Q., S.C., C.J.M., A.P.D., J.S.

Supervision: M.M.W., A.M.P., J.S.

Project Administration: J.S.

Funding Acquisition: J.S.

## METHODS

### Ethics statement

Tumor samples were collected from a SCLC patient under approval of the Institutional Review Board at Stanford University (IRB protocol #45112). Mice were maintained according to practices prescribed by the NIH, the Institutional Animal Care and Use Committee (IACUC) at Stanford University, and Association for Assessment and Accreditation of Laboratory Animal Care (AAALAC). The study protocol was approved by the Administrative Panel on Laboratory Animal Care (APLAC) at Stanford University (protocol #APLAC-13565).

### SCLC cell lines and cell culture

N2N1G cells were derived from a lymph node metastasis in an *Rb/p53/p130* mutant mouse. 16T cells derived from a late-stage primary tumor in an *Rb/p53* mutant mouse. In experiments where 16T cells needed to be visualized, they were infected with a lentivirus expressing GFP (Yang et al., 2019). KP22 cells were derived from a primary lung tumor in an *Rb/p53* mutant mouse. 5PFBl cells were derived from pleural fluid cancer cells from an *Rb/p53/p130* mutant mouse. Human SCLC cell lines were purchased through ATCC, except NJH29, which was derived from a patient-derived xenograft (PDX) as described before (Jahchan et al., 2013), and SUBr1, which was generated from a brain metastasis in a SCLC patient at Stanford University. Briefly, upon dissection and mincing using a razor blade, the samples were digested using collagenase and dispase with agitation at 200 rpm at 37°C for 15 min. The samples were treated with DNAse1 and filtered through a 70 µm membrane before expansion in recipient mice. Histological analysis (by C.K.) of xenografts confirmed features of SCLC. RNA-seq analysis (GEO number GSE178743) shows that this SUBr1 cell line belongs to the SCLC-A subtype and has high levels of NFIB.

Cells were cultured for in RPMI + 10% FBS + 1% Pen/Strep/Glutamine at 37°C with 5% CO_2_. For experiments, healthy cells were harvested from cultures with > 85% viability as determined by cell counting with trypan blue. To dissociate aggregates into single cells, the cells were pelleted, trypsinized with 0.05% Trypsin-EDTA for 1 min, gently filtered through a 40 µm filter.

### Co-culture assays with SCLC cells and astrocytes

Human and mouse astrocytes were obtained from ScienCell Research Laboratories (#1800, #M1800) and cultured in specific medium (AM #1801, a-AM #1831). For co-culture, Transwell plates with 0.4 µm pore size were used (Corning). SCLC cells were cultured in the top well (non-adherent), astrocytes were cultured on the bottom well (adherent) in medium with 1:1 mix of RPMI and astrocyte medium (AM) + 1% Pen-Strep/Glutamine + 1% FBS (RPMI-AM). Astrocytes were plated 24 hrs before adding SCLC cells into top wells. For culture with only SCLC cells, these cells were cultured in the top well in RPMI-AM. For astrocytes-only conditions, astrocytes were cultured on the bottom well in RPMI-AM.

For treatment with Tiplaxtinin and recombinant proteins (human and mouse Reelin, human SERPINE1), cells were cultured in RPMI and astrocyte medium (AM) + 1% Pen-Strep/Glutamine + 1% FBS (RPMI-AM) for co-cultures for 24 hours before adding 5µM Tiplaxtinin (SelleckChem, #S7922) or DMSO control. 100ng/mL recombinant human Reelin (R&D Systems, #8546-MR-050) and mouse Reelin (R&D Systems, #3820-MR), human SERPINE1(R&D Systems, #1786-PI) were used in the *in vitro* culture.

### Chemotaxis assays

Astrocyte chemotaxis was assessed in the Transwell migration system with membranes with 8 µm pore size (BioVision, #K906). Astrocytes were cultured in the top well for 4 hrs before adding SCLC conditioned medium or medium with recombinant proteins/antibodies/inhibitors into the bottom well. Migrated cells were lysed and detected 48 hrs later using a BioTek plate reader with Gen5 software.

### Cell viability and cell death assays

Cell viability was assessed by standard AlamarBlue assay. 10 µL of AlamarBlue reagent were added to 100 µL of cells in 96-well plates and incubated at 37°C with 5% CO_2_. Fluorescence was measured after 3 hrs of incubation using a BioTek plate reader with Gen5 software (excitation wavelength: 530nm, emission wavelength: 590nm).

Apoptosis of cells cultured *in vitro* were assessed using a cleaved caspase3/7 activity kit following the manufacturer’s instructions (Promega, cat#8090). Briefly, 100 µL of detection reagents were added to 100 µL of serum-free RPMI containing 10,000 cells in 96-well plates and incubated at room temperature for 1 hr. Luminescence intensity was measured using a BioTek plate reader with Gen5 software.

### Human cortical spheroids and fusion with SCLC

Brain region-specific human cortical spheroids (hCSs) were differentiated from two human iPSC line grown in feeder-free condition, according to the protocol described before (Yoon et al., 2019). In brief, cortical organoids were generated using Aggrewell 800 (Stem Cell Technologies), and single cell suspension of iPSCs at a density of 3×10^6^ cells/well. Dorsal forebrain region specificity was achieved by addition of dorsomorphin (5 μM, Millipore Sigma, P5499), SB-431542 (10 μM, Tocris, 1614) and XAV-939 (0.5 μM, Tocris, 3748) to Essential 6 medium (Thermo Fisher Scientific, A1516401) for the first five days. On the sixth day in suspension, neural spheroids were transferred to neural medium containing Neurobasal A (Thermo Fisher Scientific, 10888022), B-27 supplement without vitamin A (Thermo Fisher Scientific,12587010) and GlutaMax (2 mM, Thermo Fisher Scientific, 35050061), and penicillin and streptomycin (100 U ml–1, Thermo Fisher Scientific, 15140122). This neural medium was supplemented with growth factors EGF (20 ng ml–1, R&D Systems, 236-EG) and FGF2 (20 ng ml–1, R&D Systems, 233-FB), and changed every day until day 15, then every-other day until day 24. To promote differentiation of the neural progenitors into neurons, the neural medium was supplemented with neurotrophic factors BDNF (20 ng ml–1, Peprotech, 450-02) and NT3 (20 ng ml–1, Peprotech, 450-03), with medium changes every other day. After day 43 cortical organoids were maintained in neural medium as described above.

At day 200, hCSs were infected with a AAV expressing mCherry under the human GFAP promoter (AAV-GFAP::mCherry) as described before (Birey et al., 2017; Sloan et al., 2018). The expression was monitored for 1-2 weeks, before fusing to SCLC spheres. SCLC aggregates were generated using Aggrewell 800 (Stem Cell Technologies). 3×10^6^ N2N1G cells were added to each well and centrifuged for 5 min at 100xg. Cells were grown in Aggrewell for 24 hrs to allow for sphere growth. Then spheres were moved to low-attachment plates and cultured for another 3 days before fusing to cortical spheroids. Assembloids were generated by adding a cortical spheroid on top of one or two SCLC aggregates in a 1.5 mL Eppendorf tube filled with 1.2 mL neural medium. Assembloids were grown in the Eppendorf tube for 5 days for live imaging and another 5 days for immunostaining analysis.

### Live imaging of human cortical organoid-SCLC fusion

For confocal live imaging, human cortical organoid-SCLC assembloids were transferred to a well of a 96–well plate (glass-bottom plates, Corning) in 200 μL of neural medium, which was changed every 24 hrs. The fusions were kept in environmentally controlled conditions (37°C, 5% CO_2_) using a Zeiss LSM980 confocal microscope with a motorized stage. Images were acquired every 20 min for 72 hrs using a 10x objective at a depth of ∼220 μm.

### Injection of cancer cells in mice

SCLC cell lines derived from murine SCLC tumors can form allografts after injection under the skin or in the liver after tail vein injection in NOD-scid IL2Rgamma^null^ mice (NSG mice) (Denny et al., 2016; Jahchan et al., 2016; Yang et al., 2019). Briefly, all procedures were performed under general anesthesia with isoflurane and care was taken to keep the surgical site clean. For stereotactic brain injections, the mouse skin was prepared with electric shaving followed by sterilization with 70% ethanol and povidine application. A small midline incision was made with a scalpel to expose the skull. A 25 g hole was bored into the parietal bone 1 to 2 mm rostral to the central suture and right of the sagittal suture. 10^5^ cells were resuspended in 10 μL PBS and injected using a rigid 32 g needle into the subcortical region at a depth of 3mm below the level of the skull at a rate of 0.1 μL per second. After allowing 2 min for pressure equilibration, the hole in the skull was closed with bone wax and the skin overlaying the skull was reapproximated using nylon sutures. For subcutaneous tumors, 10^6^ cells were suspended in 100 μL PBS, mixed with Matrigel 1:1 and directly injected under the skin in the flank. For the intravenous injections and liver metastases, 10^6^ cells in 100 μL PBS were injected directly into the tail vein. Similar procedures were used for human cells

### Preparation of single cell isolation from mouse brain tumors

To prepare a suspension of single cells, brain tumors were harvested from mice at 3 weeks post-transplant, mechanically disrupted using a razor and digested using collagenase and dispase with agitation at 200 rpm at 37°C for 5 min. The samples were treated with DNAse1 and filtered through a 40 µm membrane and stained with DAPI. GFP-positive, DAPI-negative cells were selected via flow cytometry with GFP-negative and boiled cells serving as two separate controls. 5,000 per sample cells were barcoded and libraries were generated using the V2 10X Chromium system. The samples were aggregated and sequenced using HiSeq 4000 with a target of 100,000 reads per cell. Sequencing data can be accessed via GEO (GEO number GSE179032).

### Single-cell RNA-sequencing analysis

We generated single-cell transcriptomes using HiSeq 4000, excluding cells with low coverage, high mitochondrial genetic content or clear immune signatures, and data from the retained cells (∼30,000) was dimensionally reduced via UMAP of principle components from the 2000 genes with the greatest variability. Clusters were annotated by unsupervised density-based clustering methods (Butler et al., 2018; Stuart et al., 2019). Reads were aligned using 10X Chromium Cell Ranger software to the mm10 (Ensembl 93) reference genome, with a modification to include the 717 bp coding sequence for the EGFP protein (Addgene plasmid 26123). The number of genes captured and percentage transcripts of mitochondrial origin, standard quality control metrics, were used to assess the quality of the cells captured. Cells with fewer than 500 unique genes, greater than 6000 unique genes or greater than 5 percent of transcripts from mitochondrial genes were excluded. Additionally, cells with no transcripts of EGFP or nonzero number of transcripts from MHC class II HLA genes were also excluded. Cell cycles were regressed and expression levels were normalized using Seurat. Only highly variable genes were used for analysis. The normalized expression data was dimensionally reduced using PCA, and the results of unsupervised clustering were visualized using UMAP and t-SNE.

To compare cells from independent replicates, alignment data from different samples were combined, and similar QC methods were applied. To identify enrichment of GO biological processes within our data, genes within each cluster that were significantly different in expression levels (alpha 0.05) and upregulated by greater than 1.5-fold were analyzed using PANTHER Overrepresentation Test. Results were adjusted for false discovery rate. To compare aggregates of cells from tumors of the same microenvironment (brain, liver or subcutis) or between aggregates of cells from different microenvironments, all data were first merged and analogous analyses were then performed.

### Tissue preparation for histology

Mouse tissues were dissected from animals immediately after euthanasia. Tissues were briefly rinsed in PBS and fixed in 10% Formalin overnight. Then, tissues were transferred to 70% EtOH before paraffin embedding. For the analysis of the brain after intracranial injection, coronal cuts were made at the coordinates of initial injection site post-fixation. 4 µm paraffin sections were obtained from each sample.

### Immunofluorescence and immunohistochemistry staining

Paraffin sections were re-hydrated by 5 min-serial immersion in Histo-Clear, 100% EtOH, 95% EtOH, 70% EtOH, and water. Antigen retrieval was performed using H-3300 Citrate-Based Antigen Unmasking Solution (Vector Laboratories) at boiling temperature for 15 min. Slides were then washed in PBST (PBS + 0.1% Tween-20) for 10 min. Samples were blocked by blocking buffer (PBST + 4% horse serum) for 1 hr at room temperature and then incubated with specific antibodies against the proteins of interest at 4°C overnight. Slides were washed in PBST (PBS + 0.1% Tween20) for 3 times (10 min each) and incubated with secondary antibodies at room temperature for 2 hrs. Slides were washed in PBST (PBS + 0.1% Tween-20) for 3 times (10 min each) and incubated with PBS + DAPI. Slides were washed in PBS for another 5 min and mounted with ProLong Gold anti-fade mounting solution (Thermo Fisher Scientific). The following antibodies were used: anti-Reelin (Abcam, ab78540), anti-GFAP (Abcam, ab7260), anti-GFP (Abcam, ab13970), anti-NFIB (Abcam, ab186738), anti-NCAM (Millipore, AB5032), anti-S100B (Abcam, ab52642), anti-Vimentin (Cell Signaling, 5741S), anti-VLDLR (Novus Biologicals, NBP1-78162), anti-SERPINE1 (Novus Biologicals, NBP1-19773), anti-MAP2 (Cell Signaling, 4542), anti-CXCL1 (Proteintech, 12335-1-AP), anti-ICAM1 (Thermo Fisher Scientific, MA5407), and anti-S100A10 (Novus Biologicals, NBP1-89370).

### Immunopanning

We followed an anti-HEPACAM immunopanning protocol (Henrik Heiland et al., 2019; Zhang et al., 2016). Mouse brain tissue or brain tumors were dissected and cut into pieces using razor blades. Samples were digested using the Papain system following the manufacturer’s instructions (Worthington Biochemical Corporation). Immunopanning plates were set up as following: 20 cm plates were coated with anti-rabbit IgG at 4°C overnight. Plates were rinsed in ice-cold PBS 3 times and then coated with anti-CD45 antibody or anti-HEPACAM antibody at 4°C overnight. Plates were then rinsed in ice-cold PBS 3 times to ensure specific binding. Dissociated samples were added to the immunopanning plate with anti-CD45 and incubated for 30 min. Unbound samples were then transferred to anti-HEPACAM plates and incubated for 30 min. Then anti-HEPACAM plates were washed with PBS for 5 times. Cells bound to the anti-HEPACAM plates were harvested after trypsinization.

### Quantitative immunoassay and immunoblot analysis

Cells were lysed in TNESV buffer (50 mM Tris-HCl pH 7.5, 1% NP40, 2 mM EDTA, 100 mM NaCl) supplemented with protease inhibitors (10 µg/mL aprotinin, 10 µg/mL leupeptin, and 1 mM PMSF). Total protein was quantified using the Pierce BCA Protein Assay Kit (ThermoFisher Scientific, #23227). For quantitative immunoassays, the capillary-based Simple Western™ assay was performed on the Wes™ system (ProteinSimple) according to the manufacturer’s protocol, with 1 μg of protein used per lane. Compass software (ProteinSimple) was used for protein quantification.

### Bulk RNA seq analysis

Transcriptomic analysis of bulk cell samples was performed on SCLC-associated astrocytes (n=3 mice) and astrocytes from the sham injection injury site (n = 3 mice) for mouse astrocytes and on NCI-H82 cultured with human astrocytes (n=3), human astrocytes cultured with NCI-H82 (n=3), and with NCI-H69 (n=3). Data can be accessed via GEO (GEO number GSE178743).

Quality of raw sequencing data assessed with FastQC. Transcript expression was quantified into pseudocounts with Salmon v.0.11.3 (Patro et al., 2017).

RNA-seq pseudocounts were normalized and underwent regularized log 2 transformation using DESeq2 package v.1.28.1 (Love et al., 2014) from Bioconductor in R-studio 1.3.1093, R v.4.0.3 (R Foundation for Statistical Computing). Gene Ontology (GO) enrichment analysis performed in R using the package clusterProfiler v.3.16.1 (Yu et al., 2012) and ShinyGo v0.66 (Ge et al., 2020).

For GSEA comparison to human gene lists from (Zamanian et al., 2012b), mouse-to-human homologs were downloaded using biomaRt package (Durinck et al., 2009) in Bioconductor, and mouse homologs of gene lists were used as query gene sets for GSEA (Subramanian et al., 2005). as implemented in clusterProfiler package v.3.18.1.

## RESOURCE AVAILABILITY

Further information and requests for resources and reagents should be directed to and will be fulfilled by the Lead Contact.

## MATERIALS AVAILABILITY

Cell lines generated in this study are available upon request to the Lead Contact.

## DATA AND CODE AVAILABILITY

The code for the RNA sequencing analyses is available at Zenodo, DOI: 10.5281/zenodo.5068366. Raw sequencing data will be available at GEO (GSE179032 and GSE178743).

