## Supplemental Figures for "Neuronal mimicry generates an ecosystem critical for brain metastatic growth of SCLC"

### FIGURE S1

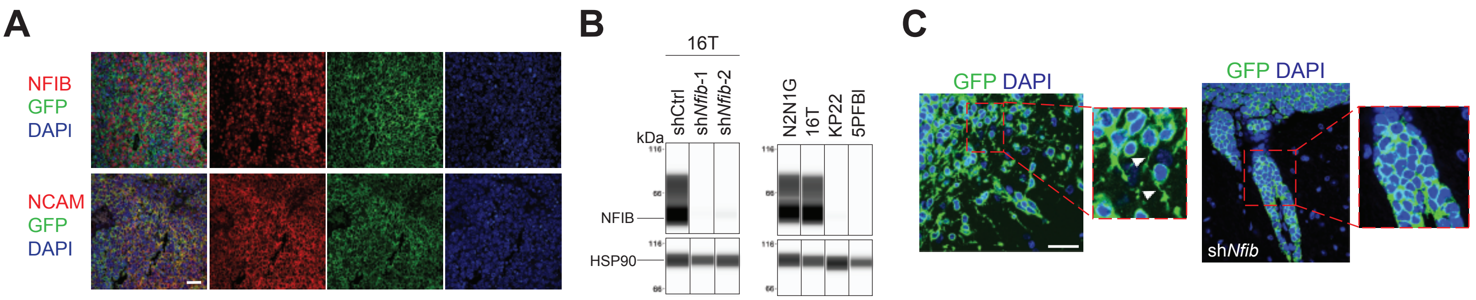


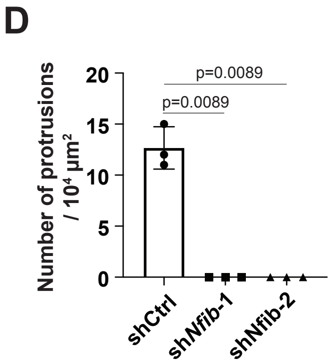


E

| Cell line | Species | Origin | NFIB status | Brain tumor+/ mice number |
|---|---|---|---|---|
| NCI-H446 | Human | Pleural effusion metastasis | High | 5/9 |
| NCI-H2081 | Human | Lung primary tumor | High | 4/9 |
| NCI-H69 | Human | Lung primary tumor | High | 1/6 |
| SUBr1 | Human | Brain metastasis | High | 5/6 |
| NJH29 | Human | Lung primary tumor | Low | 3/9 |
| NCI-H524 | Human | Lymph node metastasis | Low | 0/9 |
| N2N1G | Mouse | Lymph node metastasis | High | 9/9 |
| 16T | Mouse | Lung primary tumor | High | 9/9 |
| KP22 | Mouse | Lung primary tumor | Low | 2/9 |
| 5PFBI | Mouse | Pleural effusion metastasis | Low | 0/9 |

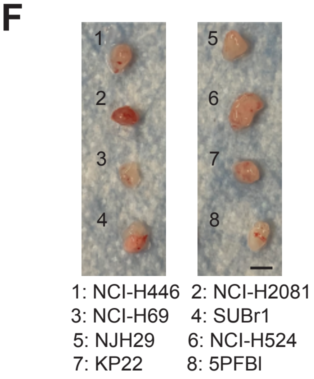


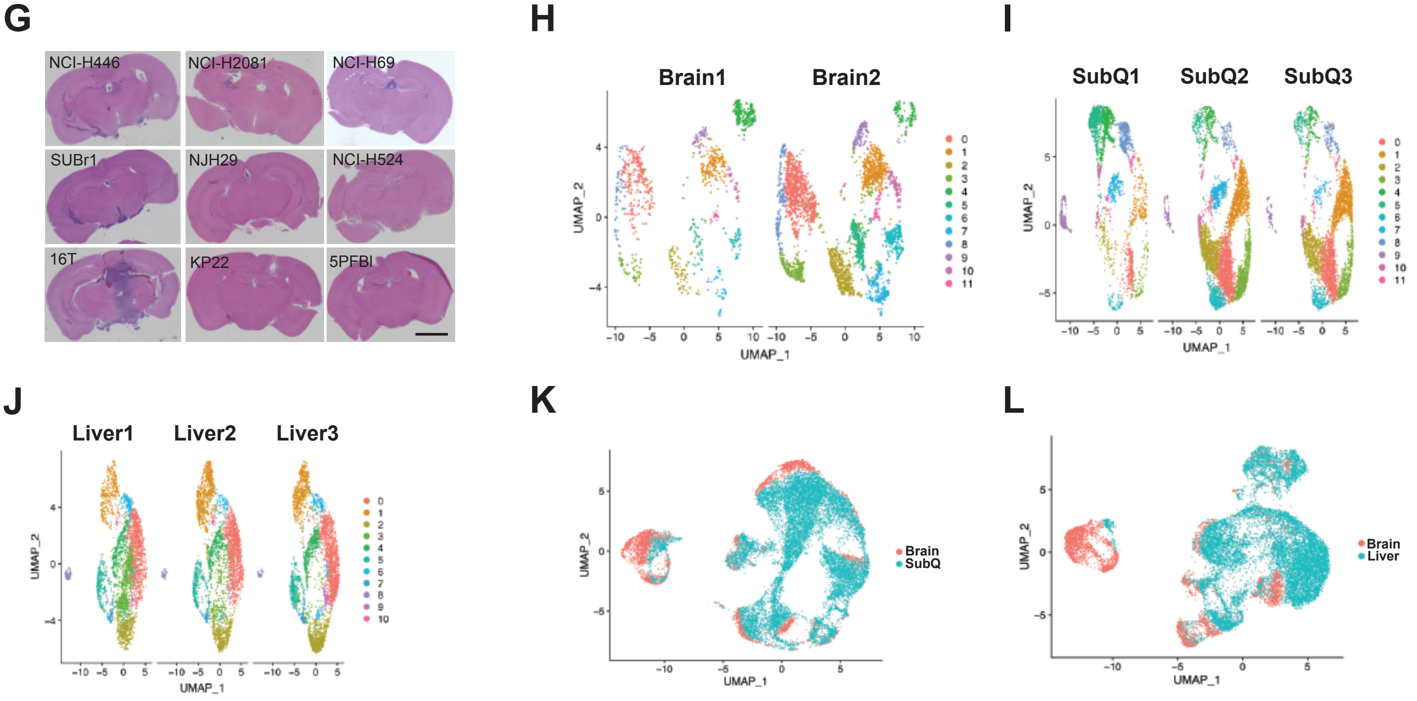


**Figure S1. Neuronal features of SCLC cells are associated with their growth in the brain**

**A.** Immunofluorescent staining for NFIB and NCAM on the sections from SCLC brain allografts as in Figure 1E. Scale bar, 20 µm. **B.** Immunoassay (by WES capillary transfer) for NFIB expression in NFIB-high 16T cells with control (Ctrl) and *Nfib* shRNAs, and in NFIB-low KP22 and 5PFBI cells. HSP90 serves as a loading control. **C**. Neuronal morphology (axon-like protrusions) in GFP-positive N2N1G cells growing in the mouse brain and loss of protrusions upon *Nfib* knock-down (sh*Nfib*). Scale bar, 20 µm. **D.** Quantification of (E) (n=3 tumors from 3 independent experiments). P values calculated via two-tailed t-test. **E.** Summary of the ability of various mouse and human SCLC tumors to grow in the brain and subcutaneously following intra-cranial injection. **F.** Representative images of subcutaneous xenografts and allografts from models in (E) after 14 days in NSG mice. Scale bar, 5mm. **G.** Representative images of brains sections from models in (E). **H-J.** UMAP clustering of 2 brain (), 3 subcutaneous (K), and 3 liver (L) single-cell RNA-seq data. **K-L.** UMAP clustering to compare single-cell RNA-seq of N2N1G cells growing in the brain vs. subcutaneous (K) and brain vs. liver (L).

### FIGURE S2

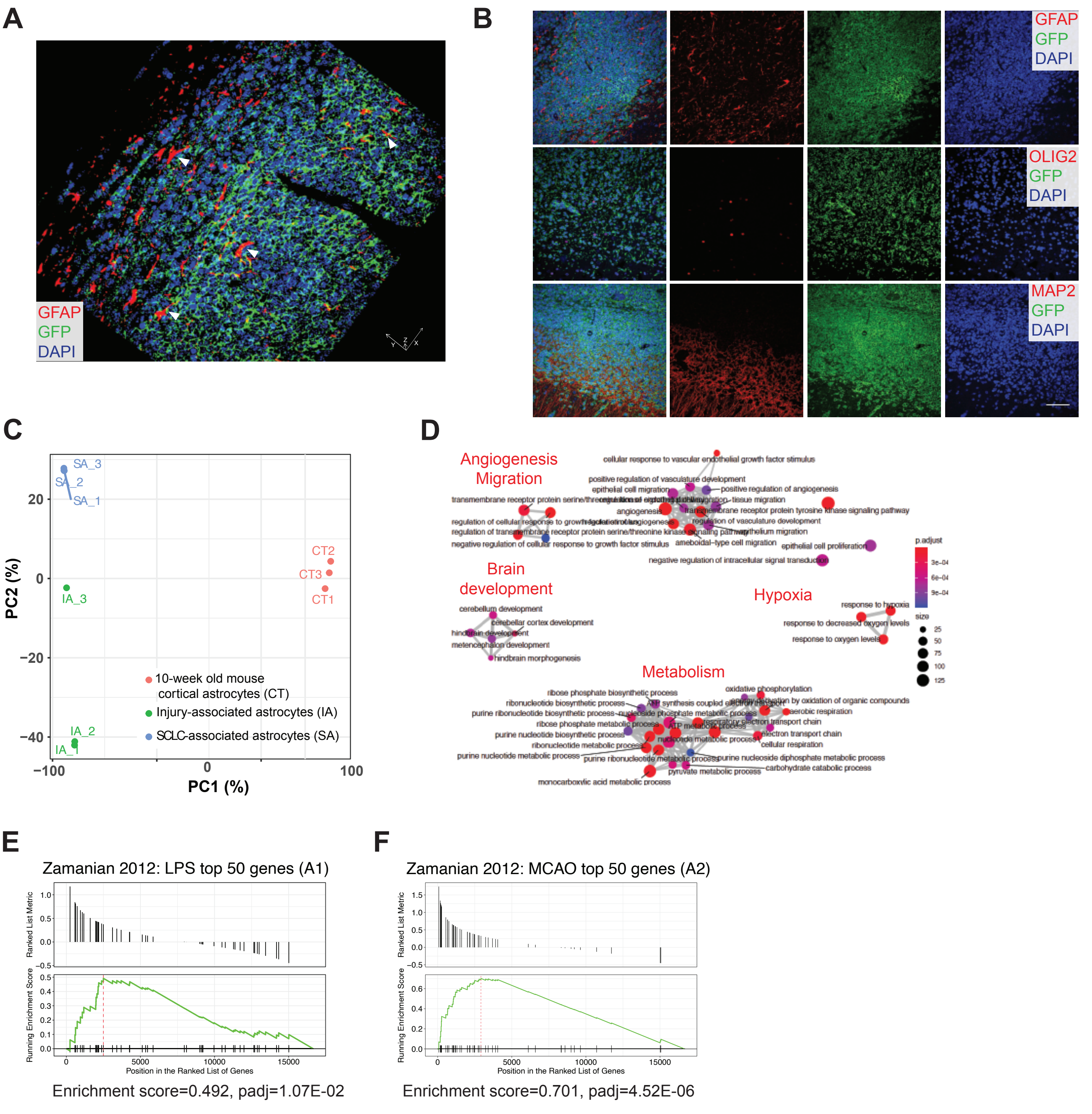


**Figure S2. GFAP-positive astrocytes infiltrate SCLC**
**A.** Representative immunofluorescent staining images of GFAP-positive astrocytes on a section from a 16T SCLC brain allograft. Arrowheads show astrocytes. Blue, DAPI to stain DNA. Scale bars: X, 50 μm, Y, 50 μm, Z, 12 μm. **B.** Representative immunofluorescent staining images of GFAP, MAP2 (mature neurons) and OLIG2 (oligodendrocytes) on a section from a 16TG SCLC brain allograft. Blue, DAPI to stain DNA. Cancer cells are GFP-positive green). Scale bar, 50μm. **C.** Principal component analysis (PCA) from the RNA-seq data comparing tumor-associated astrocytes, injury-activated astrocytes (non-tumor), and wild-type control astrocytes (n=3 each). **D**. Gene ontology (GO) enrichment network for the genes that are upregulated in SCLC-associated astrocytes compared to injury-associated astrocytes. **E-F**. Gene Set Enrichment Analysis (GSEA) for the genes upregulated in the SCLC-associated astrocytes compared to cortical astrocytes from 10-week-old mice for the top 50 genes characterizing A1 neurotoxic (E) and A2 neuroprotective (F) reactivated astrocytes.

#### FIGURE S3

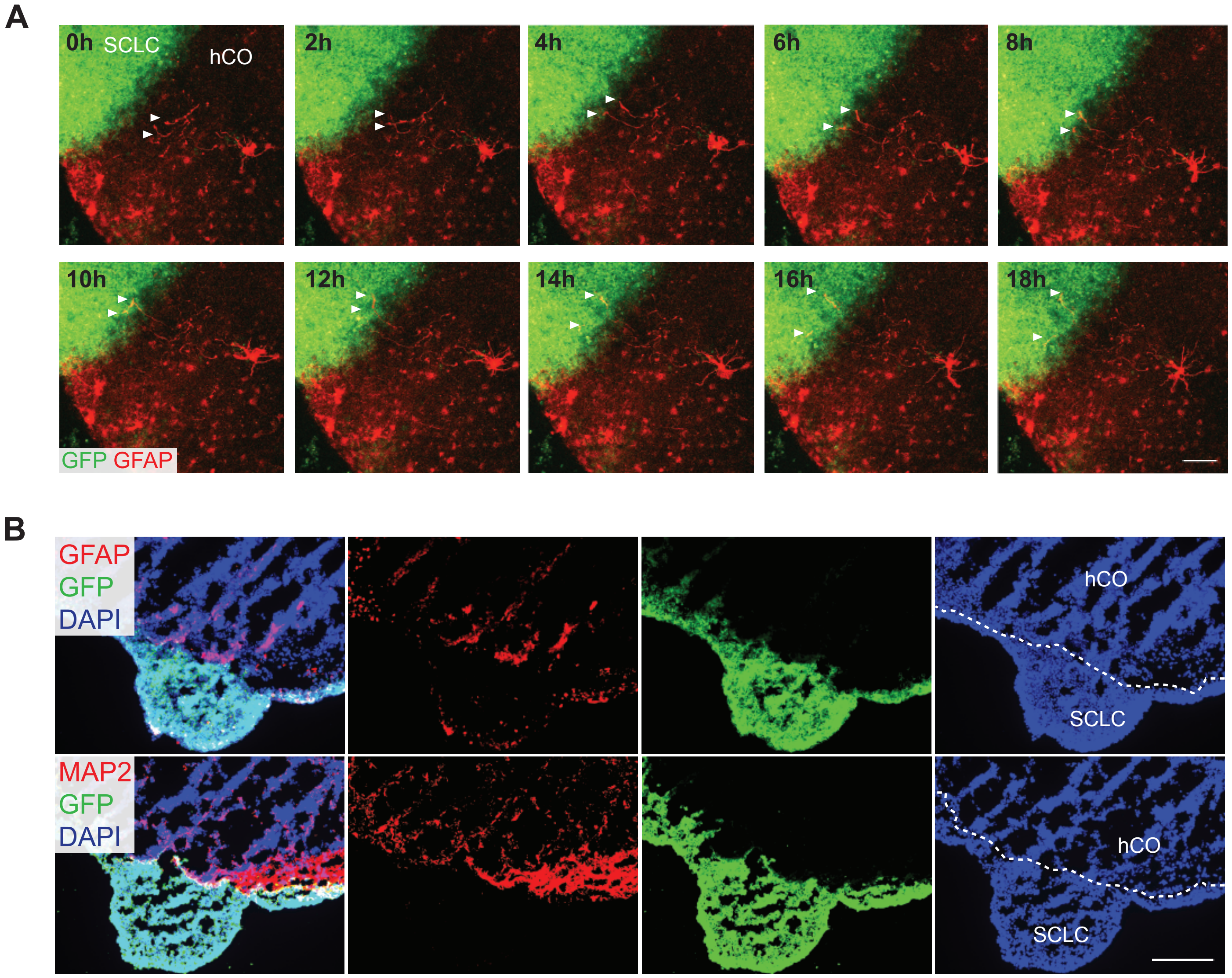


**Figure S3. Human GFAP-positive astrocytes infiltrate SCLC**

**A.** Representative images time-series from SCLC-hCO co-cultures showing the astrocyte process movement toward SCLC aggregate at post-fusion day 5-6. Red: GFAP, astrocytes. Green: GFP, SCLC. Scale bar, 50µm. Same images as in Fig. 3D but additional time points are shown **B**. Representative images showing immunofluorescent staining of GFAP (red), MAP2 (red), and GFP (green for SCLC) on sections from SCLC-hCO co-cultures at post-fusion day 10. Blue: DAPI. Scale bar, 100 µm.

### FIGURE S4

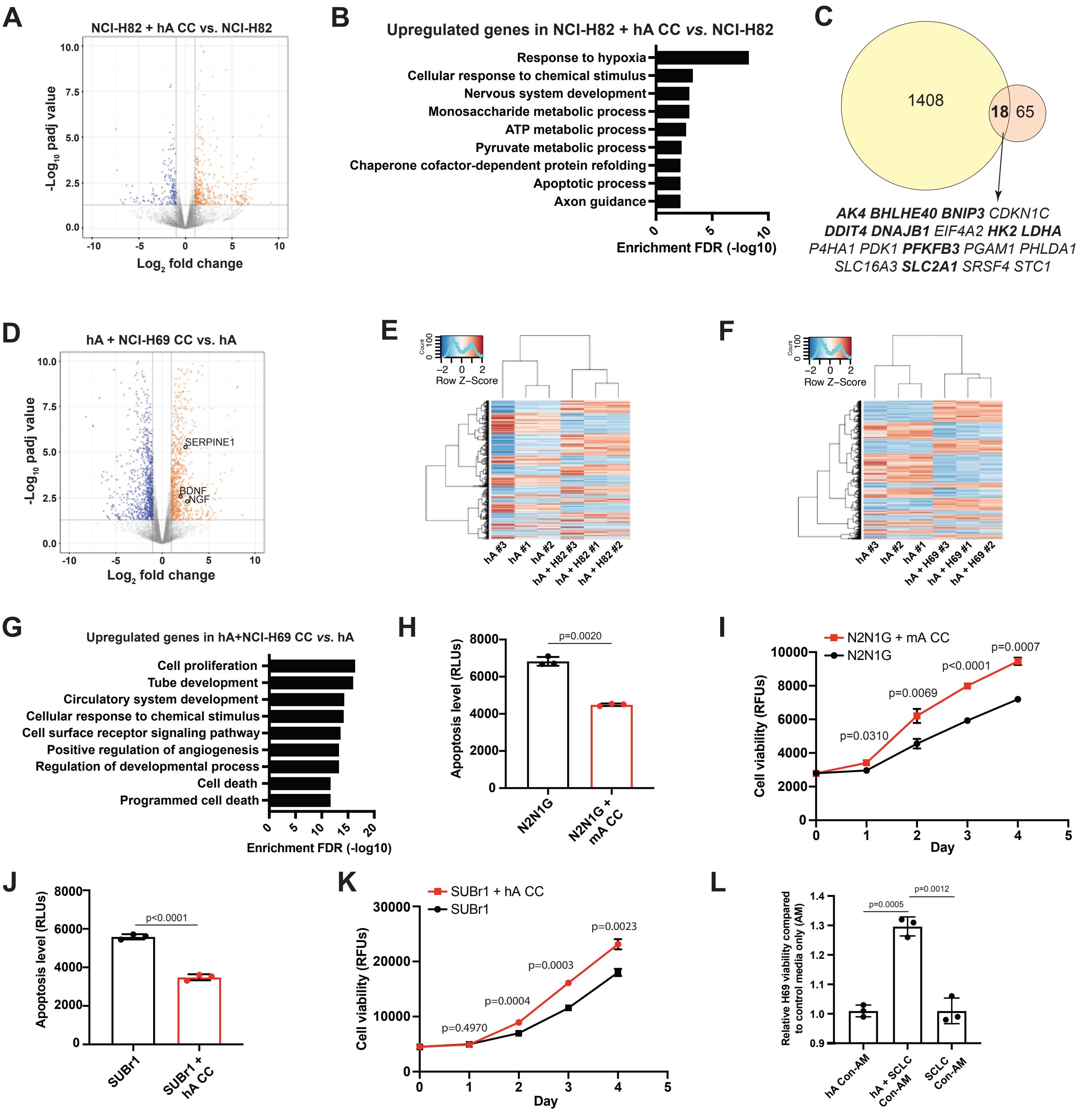


**Figure S4. Astrocytes activated by SCLC cells secrete factors promoting the survival of SCLC cells**
**A.** Volcano plot showing differentially expressed genes in NCI-H82 co-cultured with hA compared to NCI-H82 alone. **B**. Gene ontology (GO) enrichment (top 10) for the genes that are upregulated in NCI-H82 cells co-cultured with human astrocytes (hA). **C**. Overlap between upregulated genes in NCI-H82 cultured with hA and N2N1G cells growing in the brain. Highlighted genes are involved in neuronal development. **D**. Volcano plot showing differentially expressed genes in hA co-cultured with NCI-H69 cells compared to hA alone. Selected genes are indicated. **E-F**. Hierarchical clustering of gene expression in hA cultured alone and hA co-cultured with NCI-H82 (H82, C) and NCI-H69 (H69, D) cells. **G**. Gene ontology (GO) enrichment (top 10) for the genes that are upregulated in hA co-cultured with NCI-H69 cells. **H-I**. Apoptosis (caspase3/7 activity) (H) and cell viability (AlamarBlue assay) (I) measured in N2N1G SCLC cells cultured with (red) or without (black) mouse astrocytes (mA) (n=3 independent experiments). **J-K**. Apoptosis (J) and cell viability (K) in human SUBr1 cells as in (H-I), with hA. **L**. Relative cell viability (AlamarBlue assay) measured in NCI-H69 cells cultured in hA-conditioned medium, SCLC-conditioned medium, and hA + SCLC-conditioned medium compared to control medium. All p values calculated by two-tailed t-test.

#### FIGURE S5

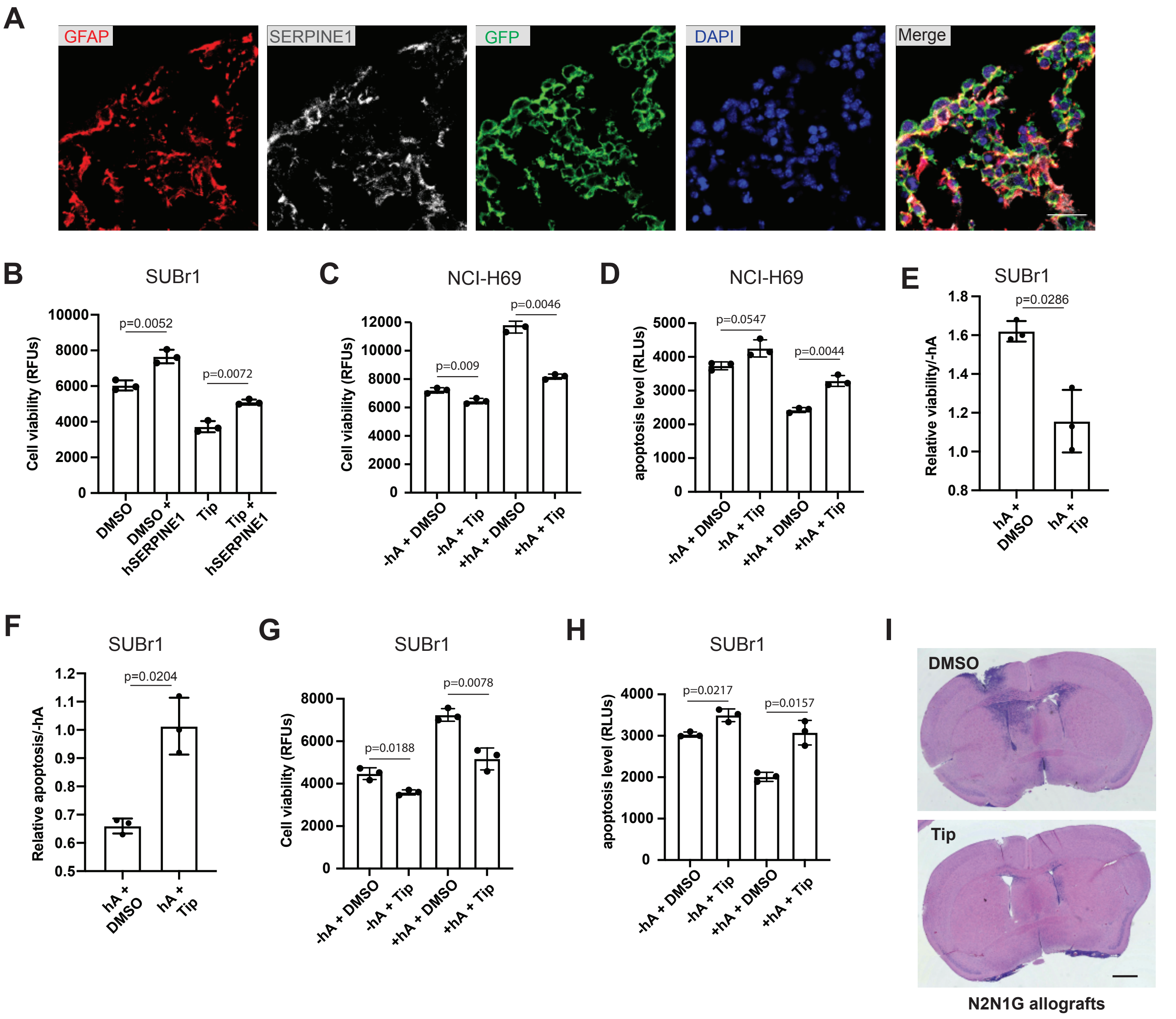


**Figure S5. SERPINE1 from astrocytes promotes SCLC growth**

**A.** Representative images showing immunofluorescent staining of GFAP (red), SERPINE1 (white), and GFP (green for SCLC) on sections from a SCLC/mini-brain at post-fusion day 10. Blue: DAPI. Scale bar, 20 μm. **B.** Cell viability (AlamarBlue assay) measured in SUBr1 cells treated with recombinant hSERPINE1 and the SERPINE1 inhibitor Tiplaxtinin (Tip) (n=3 independent experiments). **C.** Cell viability (Alamar-Blue assay) measured in NCI-H69 cells cultured with human astrocytes (hA) or without hA, with or without Tip treatment (n=3 independent experiments). **D.** Apoptosis (caspase3/7 activity) measured in NCI-H69 cells cultured with hA compared to without hA, with or without Tip treatment (n=3 independent experiments). **E.** Cell viability (AlamarBlue assay) measured in SUBr1 cells cultured with hA compared to without hA, with or without Tip treatment (n=3 independent experiments). **F.** Apoptosis (caspase3/7 activity) measured in SUBr1 cells cultured with hA compared to without hA, with or without Tip treatment (n=3 independent experiments). **G-H**. Same as (C-D) for SUBr1 cells and human astrocytes. **I.** Representative images of hematoxylin and eosin-stained (H&E) coronal sections from the brain of recipient mice following injection of mouse N2N1G SCLC cells at day 14. Cells were incubated with DMSO or the SERPINE1 inhibitor Tip at the time of injection. Scale bar, 1 mm. All p values calculated by two-tailed t-test.

#### FIGURE S6

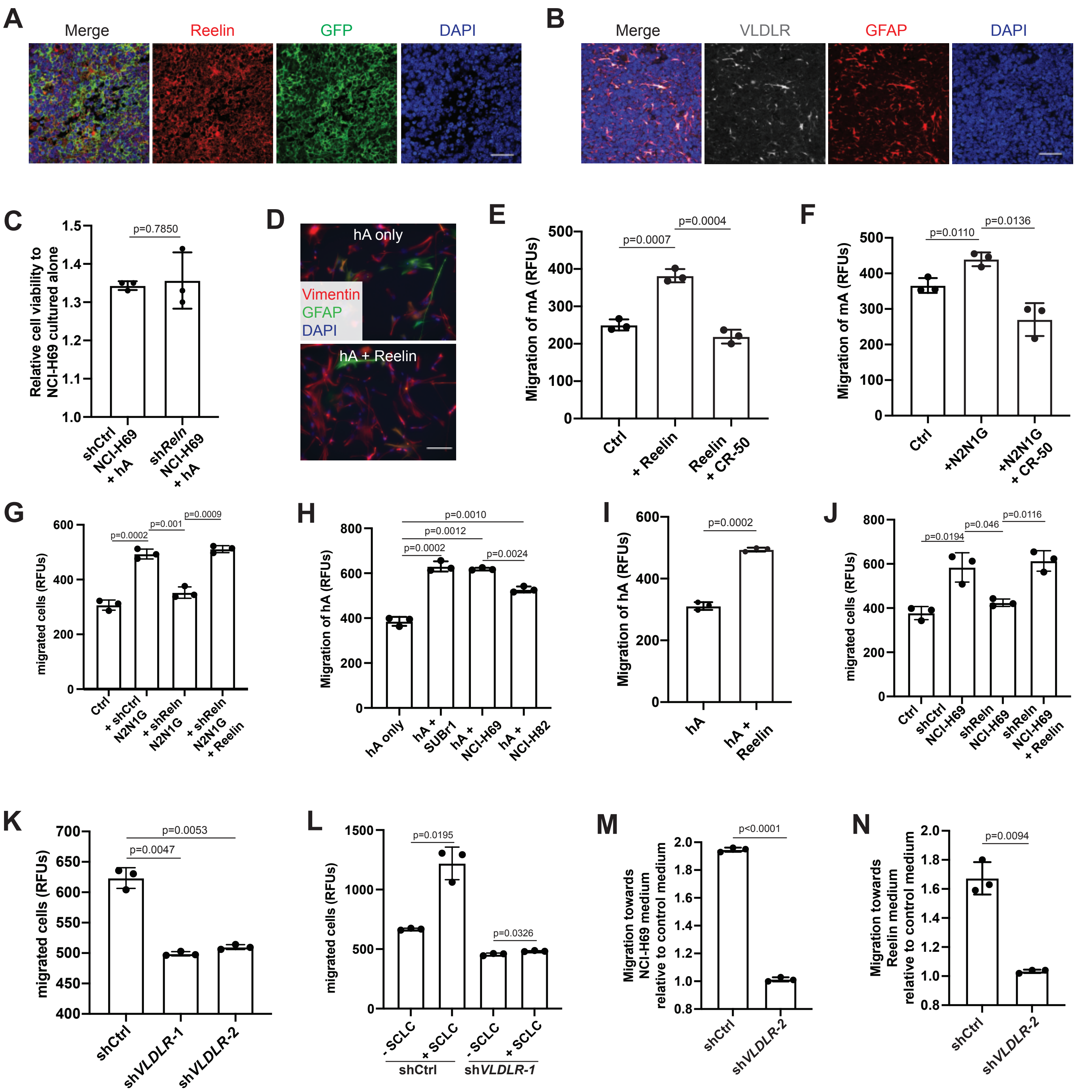


**Figure S6. Reelin-VLDLR signaling regulates SCLC-induced astrocyte migration**
**A-B**. Representative images showing immunofluorescence staining of Reelin (red) and GFP (green, cancer cells) (A), VLDLR (gray) and GFAP (red) (B) on an N2N1G brain allograft section. Blue: DAPI to stain DNA. Scale bar, 50 μm. **C**. Cell viability (AlamarBlue assay) measured in shCtrl and sh*Reln* NCI-H69 cells cultured with human astrocytes compared to without astrocytes (n=3). **D**. Representative images showing immunofluorescent staining of GFAP (green) and Vimentin (red) in human astrocytes (hA) treated with or without recombinant human Reelin. Blue: DAPI to stain DNA. Scale bar, 20 μm. **E**. Mouse astrocyte (mA) chemotaxis in control medium, with recombinant mouse Reelin or the Reelin blocking antibody CR-50 (n=3). **F-G**. Mouse astrocyte (mA) chemotaxis in control medium, N2N1G-conditioned medium, with or without the Reelin blocking antibody CR-50 (n=3) (F) and shCtrl or shReln N2N1G-conditioned medium, with or without mouse Reelin. **H**. Human astrocyte chemotaxis in control medium or hSCLC-conditioned medium (NCI-H69, SUBr1, and NCI-H82) (n=3). **I**. Human astrocyte chemotaxis in control medium and in the presence of recombinant human Reelin (n=3). **J**. Same as (G) for human astrocytes and human Reelin. **K-L**. Chemotaxis of shCtrl and sh*VLDLR* human astrocytes (K) with or without hSCLC-conditioned medium (L). (n=3). **M-N**. Chemotaxis of shControl and sh*VLDLR*-2 hA in NCI-H69-conditioned medium (M) and medium containing human Reelin protein (N) compared to control medium (n=3 ). All p values are calculated via two-tailed t-test.

### FIGURE S7

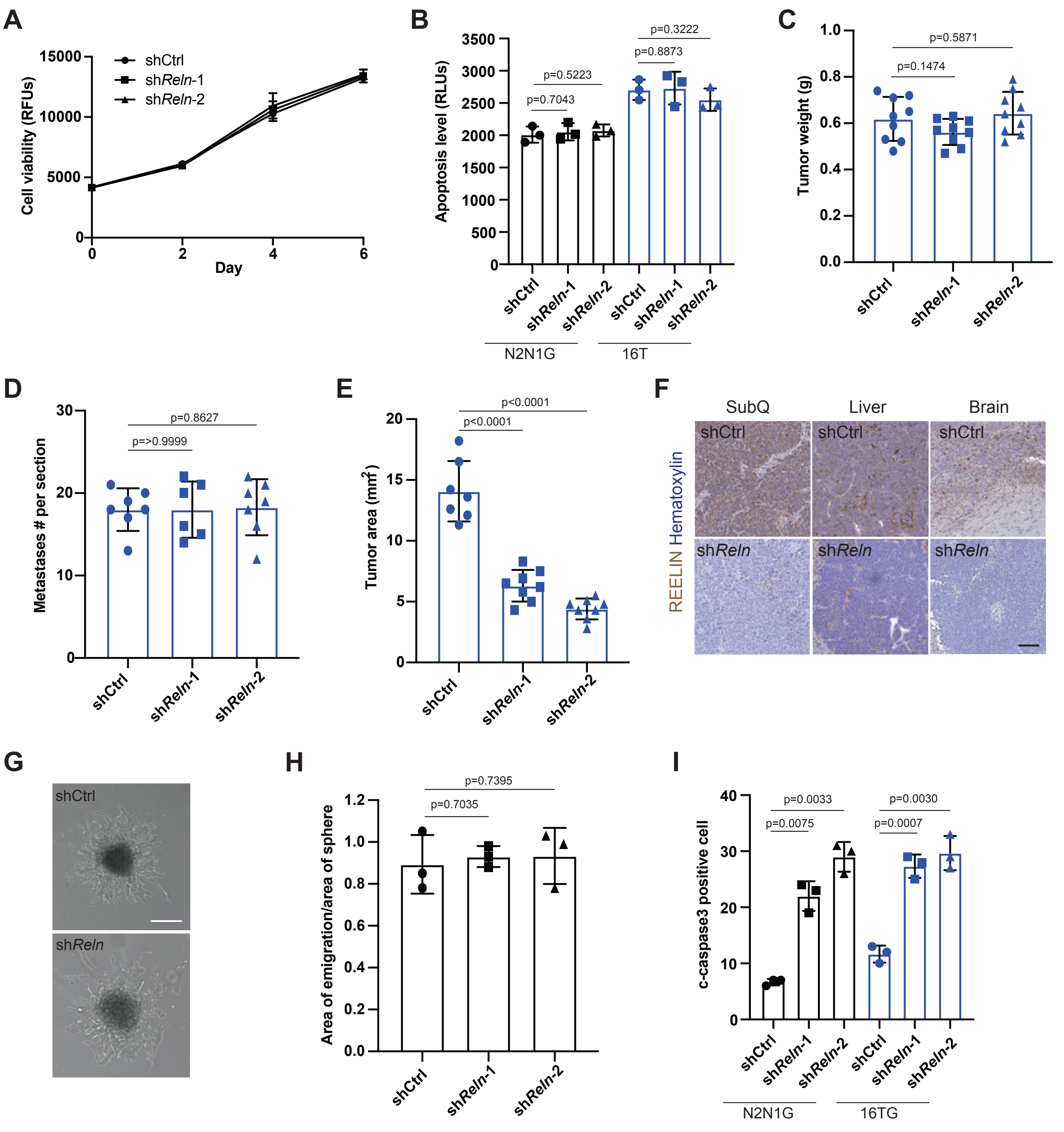


**Figure S7. Reelin expression is critical for the growth of SCLC brain tumors *in vivo***

**A.** Cell viability measured by AlamarBlue assay of shControl and sh*Reln* N2N1G cells. **B**. Apoptosis level of shControl and sh*Reln* N2N1G (black) and 16T (blue) cells measured by caspase3/7 activity. **C**. Size of subcutaneous tumors generated from shControl and sh*Reln* 16T cells. N=7, 8 from 2 independent experiments. **D**. Number of liver metastases generated from i.v. injection of shControl and sh*Reln* 16T cells. N=6, 7 from 2 independent experiments. **E**. Tumor size of shControl and sh*Reln* 16T brain allografts. N=7,8 from 2 independent experiments. **F**. Representative images of IHC staining of Reelin on shControl and sh*Reln* N2N1G subcutaneous (SubQ) tumors, liver metastases, and brain allografts. Scale bar, 100μm. **G**. Representative images of shControl and sh*Reln* N2N1G spheres growing in the 3D Matrigel culture. Day 2 after seeding. Scale bar, 50μm. **H**. Quantification of the area of emigration out from shControl and sh*Reln* N2N1G spheres. **I**. Quantification of cleaved c-caspase3 (CC3) positive SCLC cells from shControl and sh*Reln* N2N1G (black) and 16TG (blue) brain allografts. All p values are calculated via two-tailed t-test.
